# Optimization of Search and Acquisition Strategies for DSSO Crosslinking Mass Spectrometry using Protein Prospector and Touchstone

**DOI:** 10.64898/2026.10.08.757733

**Authors:** Michael J. Trnka, Clinton Yu, Peter R. Baker, Philip T. McGilvray, Fenglong Jiao, Robert J. Keenan, A.L. Burlingame, Robert J. Chalkley, Lan Huang

**Author notes:** Corresponding authors: Michael Trnka and Lan Huang. Current address: Anagram Therapeutics, Natick, Massachusetts, USA.

## Abstract

Crosslinking mass spectrometry (XL-MS) enables residue-level characterization of protein-protein interactions (PPIs) in both purified assemblies and complex biological systems. Cleavable crosslinkers such as DSSO are widely used to facilitate analysis of complex samples, yet there remains limited consensus on how best to leverage their properties in both acquisition and data analysis workflows. In part, this reflects differences in scoring, classification, and false discovery rate (FDR) estimation across analysis pipelines. Therefore, an integrated XL-MS data analysis pipeline that enables confident identification of MS-cleavable cross-linked peptides through improved scoring and rigorous FDR assessment is essential for systematically comparing and benchmarking workflows across different sample preparations. To this end, we implemented a new capability within the Protein Prospector search engine to improve MS^2^-based identification of MS-cleavable cross-linked peptides. Next, we developed Touchstone, a framework for re-scoring, classification, FDR-controlled summarization, and visualization of Prospector XL-MS search results. Using this approach, we evaluate the contributions of DSSO-specific fragmentation features and compare multiple acquisition strategies across datasets ranging from purified macromolecular assemblies to proteome-scale samples. We find that the primary advantage of DSSO arises from enhanced backbone fragmentation of both linked peptides, rather than from diagnostic ion formation, and that incorporating diagnostic ion pairs into MS2-based scoring or filtering does not improve identification performance. Benchmarking against established datasets shows that Prospector-Touchstone performs comparably to leading methods for non-cleavable crosslinkers and provides improved performance for DSSO analysis. We further compare acquisition strategies and find that for purified protein assemblies, MS2.stHCD characterizes more residue-pairs than ETD-based, MS3-based, and hybrid methods. In contrast, for proteome-scale XL-MS, MS3-based acquisition yields similar numbers of inter-protein PPIs with lower error rates despite reduced sampling depth.

Together, these results highlight complementary strengths of MS2-and MS3-based workflows and provide practical guidance for MS-cleavable XL-MS experiments with varying sample complexities.

**Highlights:**

- Prospector-Touchstone improves crosslink identification and error estimation for both cleavable and non-cleavable XL-MS workflows relative to existing analysis pipelines.
- The cleavable crosslinking reagent DSSO enhances the identification of peptide backbone ions from both linked peptides compared to non-cleavable crosslinkers.
- Incorporating DSSO diagnostic ion pairs into MS2-based scoring or filtering does not improve XL-MS identification performance.
- For purified protein assemblies, MS2.stHCD outperforms ETD-based, MS3-based acquisition, and hybrid acquisition strategies.
- For proteome-scale XL-MS, MS2.stHCD identifies more residue-pairs, while MS3.CID yields higher-confidence protein-protein interactions.

## Introduction

Crosslinking Mass Spectrometry (XL-MS) utilizes bifunctional reagents to covalently label macromolecular systems and infer spatial proximity by identifying the adducted residues with bottom-up proteomics techniques(1, 2). Over the last decade, XL-MS has matured into a robust technique capable of streamlined analysis of purified protein assemblies containing dozens to several hundred subunits. Crosslinking data of protein complexes have been used to support structural inference either by validating or challenging models built using other methods(3), or by using crosslinked distance restraints alongside other biophysical and biochemical data to guide the construction of integrative models(4). Despite recent advances in cryo-EM (5) and deep learning-based structure prediction(6, 7), XL-MS continues to provide complementary information, particularly for positioning structured domains within large, flexible assemblies, and characterizing conformationally heterogeneous complexes, such as “fuzzy”(8) or dynamic(9) complexes, as well as biomolecular condensates(10, 11).

XL-MS technology is increasingly applied to structural systems biology, where it captures endogenous protein-protein interactions (PPIs) in their native environment without cellular engineering. XL-MS enables simultaneous identification of direct PPIs and residue-level contacts from proteomes and subproteomes including isolated organelles(12, 13), whole-cell lysates(14, 15), intact cells(16, 17), and tissues(18).

However, system-scale XL-MS presents substantial analytical challenges. Database searches are performed against proteomes containing up to ∼20,000 sequences, orders of magnitude larger than those used for purified systems, and best practices for error estimation and control at this scale are still evolving(19–23). A central challenge arises from the quadratic combinatorial expansion of candidate peptide pairs (the “n-square problem”), which inflates the search space relative to linear proteomics analyses(24).

Error rates can also increase at downstream levels of data summarization when crosslinked spectral matches (CSMs) are collapsed into residue pairs and protein pairs, making it important to estimate FDR at the level ultimately being reported and to treat intra-and inter-protein assignments separately. For example, an orthogonally benchmarked proteome-scale XL-MS study found that applying a 5% CSM-level FDR and combining intra-and inter-protein assignments resulted in an estimated 35% error rate at the PPI level(20). In addition, detection of crosslinked peptides from low-abundance and transient PPIs in complex mixtures remains a major limitation, necessitating new developments in sample preparation and handling strategies(25).

MS-cleavable crosslinkers have been developed to address some of these challenges by introducing low-energy, gas-phase labile bonds that produce diagnostic ions corresponding to the individual peptides(1, 26). Various functionalities have been used to generate the labile site including an Asp-Pro bond(27), a urea moiety(28), or a Rink linker(29). One of the most widely used cleavable reagents is DSSO(30), which utilizes a sulfoxide group to provide gas-phase lability. In collisional dissociation, these bonds fragment to form diagnostic product ions with the mass of the linear peptides plus a defined tag. Early workflows for DSSO relied on MS3-based acquisition, in which diagnostic ions observed in MS2 are selected for further fragmentation. Searching these MS3 spectra bypasses the ‘n-square problem’ of trying to identify pairs of peptides in a single spectrum. While effective, these approaches reduce sampling efficiency due to increased cycle time and depend on accurate real-time targeting of diagnostic ions, which can be limited by low signal intensity and non-specific selection of MS3 precursors.

To improve acquisition speed and sensitivity, strategies have been developed that extract both diagnostic-ion and backbone-fragmentation information from only MS2-spectra. Some of these approaches rely on detection of diagnostic ion pairs to infer individual peptide masses prior to sequence identification, which linearizes the database search by enabling independent identification of both component peptides. Software tools such as XlinkX(31) and MeroX(32) implement this strategy and often combine multiple fragmentation modes to balance diagnostic ion formation and backbone fragmentation. For instance, a resonant CID scan to optimize yield of diagnostic ions is combined with either a second beam-type collisional dissociation (HCD) scan(33) or an electron transfer dissociation (ETD/EThcD) scan(31) to maximize backbone ion yield.

Alternately, stepped HCD (stHCD) methods(34) that employ multiple different collisional energies have been suggested to optimize both types of fragmentation pathways in a single scan. Some labs have combined MS2 and MS3 diagnostic-centered methods by using MS2-CID-MS3-CID-MS2-EThcD acquisition cycles together with search engines such as XlinkX that integrate information from the different scan types(15).

An alternative approach is provided by mass modification or open-search algorithms, including Protein Prospector(35), pLink2(36), Kojak(37, 38), and the XiSearch-XiFDR pipeline(39), which identify crosslinked peptides without the need for diagnostic ions.

These methods treat crosslinks as linear peptides bearing discrete mass modifications(40, 41). The list of top hits to a spectrum is then examined pairwise to look for complementary peptides and mass modifications consistent with the precursor ion(35). This strategy linearizes the initial peptide search and substantially reduces the combinatorial burden relative to exhaustive pairwise searching, enabling proteome-scale analysis(17, 42). These approaches were originally developed for non-cleavable crosslinkers, and their application to cleavable reagents such as DSSO has not been systematically explored. In addition, error rate estimation in MS2-based XL-MS analysis is more difficult than in MS3-based approaches(43), necessitating robust strategies for scoring, classification, and false discovery rate (FDR) estimation.

Protein Prospector incorporates a mass-modification search strategy for XL-MS, but tools for downstream interpretation of crosslink search results have been limited. To improve the utility of Protein Prospector across both non-cleavable and MS-cleavable crosslinkers, we developed Touchstone, an R-based module for rescoring, classification, and FDR-controlled summarization of Prospector XL-MS results.

Touchstone summarizes crosslinking data at multiple levels, including crosslink spectral matches (CSMs), unique residue pairs (URPs), and protein pairs, and additionally supports module annotation, visualization, and integration with structural models and STRING-db for validation.

We first benchmark Prospector-Touchstone on established non-cleavable(44) and cleavable datasets(45). Because mass-modification searches do not require diagnostic ions, this framework enables direct evaluation of the contribution of DSSO-specific fragmentation to identification performance. We then compare multiple acquisition strategies across purified assemblies and proteome-scale samples. Our results show that the primary advantage of DSSO arises from enhanced backbone fragmentation rather than diagnostic ion formation, and that incorporating diagnostic ion pairs into MS2-based scoring or filtering does not improve identification performance. We further demonstrate that optimal acquisition strategies depend on sample complexity: MS2.stHCD provides superior residue-pair identification for purified assemblies, whereas MS3-based approaches identify fewer residue-pairs but yield lower error rates while recovering a similar number of protein-protein interactions in complex systems.

Together, these results provide practical guidance for DSSO-based XL-MS workflows and clarify the roles of MS2-and MS3-based strategies in different experimental contexts.

## RESULTS

### Development of the Prospector-Touchstone Cross-link Analysis Pipeline

To facilitate automated scoring, classification, summarization, visualization and biological interpretation of Prospector XL-MS search results we developed Touchstone, an R package supporting analysis of both cleavable and non-cleavable crosslinking datasets. Touchstone can be run interactively or in scripted R environments and is also available through a Shiny web interface integrated with the Prospector pipeline workflow. The Shiny interface additionally provides tools for visualization of protein networks and inspection of annotated spectra (Supplemental Figure 1). Touchstone imports Prospector Search Compare results, constructs support vector machine (SVM)-based scoring models, and summarizes identifications at user-defined FDRs and levels of summarization, including CSMs, URPs, peptide pairs, protein pairs, and module pairs. Additional functionality includes manual spectrum review, module/domain annotation, structural mapping onto PDB models with observed and randomized Cα–Cα distance calculations, retrieval of STRING-db interaction scores, and visualization through internal plotting functions or export to xiView(46) and ChimeraX(47) pseudobond files.

Touchstone employs SVM-based classification of spectral matches using features describing precursor and product-ion match quality together with features derived from higher levels of summarization. The classifier supports independent FDR estimation at multiple summarization levels and separate thresholding of intra-and inter-protein crosslinks. Treating intra-and inter-protein crosslinks identically can substantially underestimate true error rates, particularly in proteome-scale searches (20, 35). Details of classifier construction and feature selection are described in the Methods.

### Benchmarking the Performance of Prospector-Touchstone for Non-Cleavable Crosslinkers

To benchmark the performance of the Prospector-Touchstone pipeline and compare it to existing MS2-based crosslink search engines, we used a dataset acquired by the Mechtler lab, composed of 95 synthetic Cas9-derived peptides that were reconstituted into 12 pools and crosslinked with non-cleavable reagent DSS(44). Peptide N-termini and C-terminal lysine residues were protected during the crosslinking reaction to ensure that crosslinks only occurred on internal lysine side chains. Ground truth is approximated by assuming that crosslinks between members of a pool are correct and all other hits are false positives.

Accurate estimation of FDR is critical to XL-MS analysis, where small spectral assignment errors can compound into overwhelming errors when the results are summarized as protein interactions(20). We first evaluated different methods of constructing decoy databases for modelling error rates. Many XL-MS experiments are performed on highly purified or reconstituted samples which contain relatively small numbers of proteins (less than 50). We reasoned that the traditional FDR estimation approach of using equally sized target and decoy databases would inaccurately model the incorrect hits, which affects both training of the scoring function as well as FDR estimation. Hence, we constructed decoy databases that were 10× larger than the target database and adjusted our FDR calculation to reflect the large size of the decoy search space (see Methods section).

The Cas9 benchmarking data were searched with Prospector against a FASTA database containing target sequences of Cas9 plus 116 common contaminants, as in the original study. Decoy sequences were constructed using one of four different randomization strategies: the standard Prospector 1× random database where the decoy version of each protein has the same amino acid composition as the target but scrambled order (“1×Scrambled”), a 1× database in which the distribution of amino acids was weighted to reflect their distribution in nature (“1×Natural”), a 10× database with a natural distribution (“10×Natural”) (48), and a 10× database with a natural amino acid frequency and peptide lengths that were normalized by fixing the position of Lys/Arg in the decoy sequences (“10×Natural+Tryptic”) (Supplemental Figure 2A). The numbers of incorrect hits to the target database with one or both peptides wrongly annotated were estimated by counting hits with one target and one decoy peptide (TD) and two decoy peptides (DD) and then adjusting for the increased size of the decoy database relative to the target (Supplemental Figure 2B). The FDRs estimated by decoy analyses were compared to ground truth FDRs at different score thresholds when classified on CSMs. The larger 10× databases better approximated the ground truth FDR measurements than the 1× databases (Figure 1A). In particular, the 10×Natural database resulted in the closest correspondence between ground truth and nominal FDR across a range of FDR values. Adjusting the peptide length distribution of this decoy database to match the tryptic peptide length of the target database (“10×Natural+Tryptic”) did not provide better performance.

**Figure 1.**
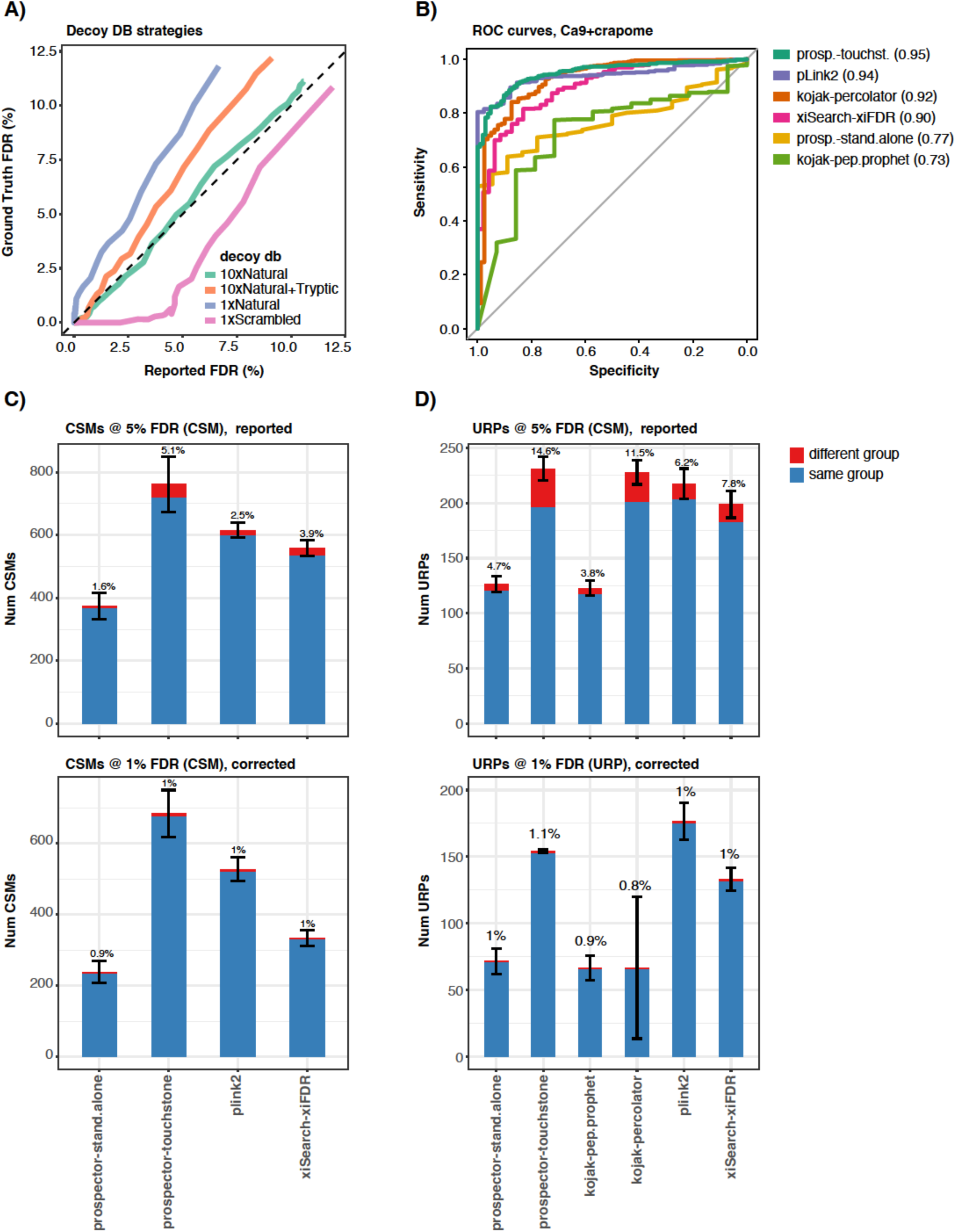
**The Prospector-Touchstone pipeline evaluated on a small crosslinking search task with a non-cleavable linker**. Searches were performed using a benchmarking library consisting of Cas9 derived synthetic peptides crosslinked with BS3 (Beveridge *et al.,* 2020) and using a database of Cas9 plus 116 abundant contaminants from the CRAPome repository. A) Evaluation of different decoy database generation strategies. The relative size of the decoy database to the target database (1x or 10x more amino acid residues) was tested as well as methods to either preserve the peptide length distribution of the target database (“tryptic”) or the natural distribution of amino acids in the proteome (“natural”). Ground truth FDR was determined by the number of disallowed crosslinks (between peptides in different groups of the benchmarking set) and the number of allowed crosslinks (between peptides in the same group) and plotted relative to reported FDR. B) Receiver operating characteristic curves for different search engines on the Cas9 plus CRAPome benchmarking library summarized on URPs. Area Under the Curve (AUC) values are reported in parentheses in the figure legend. C, D) Different mass modification based crosslinking search platforms benchmarked against the Cas9 plus CRAPome BS3 library. Average and standard deviation of three independent replicates. The experimental design replicates the methodology of Beveridge *et al* (2020) and results from other search engines are taken from the supplemental materials of that reference. Prospector – Touchstone analysis utilized a 10x scaled decoy database and a single threshold for all crosslinks. Prospector – Stand Alone analysis utilized a 1x scrambled database. C) Top panel-Search results were reported at 5% CSM FDR and the number of correct and incorrect spectral matches are plotted with the ground truth FDR reported above the bars. Bottom panel-Search results reported at a corrected 1% ground truth FDR. D) Top panel-The same analysis as C reported as unique-residue pairs (URPs). Bottom panel-Search results reported at a corrected 1% ground truth FDR determined on URPs.

We next evaluated the performance of Prospector-Touchstone against three commonly used MS2-based search engines (*i.e.* pLink2, XiSearch, and Kojak) included in the original benchmark study(44), as well as stand-alone Prospector. To enable direct comparison, we followed the same analytical workflow and integrated the Prospector results with the previously published results from the other search engines. The Cas9 benchmark library was searched using different search engines and data were classified at a 5% CSM FDR threshold. The number of correct and incorrect hits at this level are reported. Additionally, the results are reported as URPs which, as expected, resulted in inflated apparent FDR since the classification took place on CSMs. In these comparisons, Prospector-Touchstone substantially outperformed stand-alone Prospector and identified more CSMs and a comparable number of residue-pairs to pLink2, Xi-XiFDR, and Kojak-Percolator (Figure 1C, 1D top panels). Because the ground truth FDRs differ considerably between the search pipelines, we repeated this analysis reporting results at a fixed 1% ground truth CSM FDR as well as a fixed 1% ground truth URP FDR (Figure 1C, 1D bottom panels). As a result, Prospector-Touchstone identified nearly three times as many CSMs as stand-alone Prospector (236 CSMs). Notably, it also identified 29% more CSMs than pLink2 (672 vs. 522), the top-performing search engine in the original benchmark study(44). When reporting at a corrected 1% URP threshold, pLink2 reported 175 URPs vs 153 for Prospector-Touchstone. Across all four comparisons, no differences in the number of results reported by Prospector-Touchstone and pLink2 were statistically significant (p-values > 0.05). ROC curves for both the crosslink and CSM level data show that Prospector-Touchstone has the highest area-under-the-curve value, although notably, for residue-pair level data, pLink2 has better sensitivity at the very highest specificity levels that are most relevant to classifying real datasets (Figure 1B, Supplemental Figure 3).

### Benchmarking the Performance of Prospector-Touchstone for Cleavable Crosslinkers

Prospector originally implemented mass modification searching to analyze data from non-cleavable reagents such as DSS (35, 41). The product ions were calculated under the assumption that the site of crosslinking would retain the mass modification intact; *i.e.* the crosslinker and second peptide would remain attached. This differs for a cleavable reagent which readily dissociates and leaves a residual fragment mass on the peptide. Hence, the theoretical product ions for cleavable reagents were only correctly calculated for fragment ions which do not contain the crosslinked site. Although cleavage between the crosslinker and one of the linked peptides is frequently observed during analysis of nominally non-cleavable reagents, producing the P, PL, and PLK ions scored by Prospector (35), this dissociation is generally incomplete compared with the nearly quantitative cleavage of DSSO under collisional activation. To enable

Prospector-Touchstone to identify MS-cleavable crosslinked peptides, we implemented new features in Prospector Batch-Tag searching to account for the unique fragmentation of DSSO and similar reagents that produce a pair of mass tags under collisional dissociation. The utility of this feature was first demonstrated by reanalyzing a DSSO crosslinked ribosome dataset using the cleavable search option instead of the non-cleavable search parameters. This resulted in a 50-100% increase in the numbers of CSM and URP identifications and significantly improved fragment ion coverage for both linked peptides (Supplemental Figure 4).

Next, we evaluated the performance of Prospector-Touchstone using a DSSO-based XL-MS benchmarking library designed by the Mechtler lab(45). As with the Cas9 library, ground truth was established using crosslinked synthetic peptides derived from 38 *E. coli* ribosomal proteins. We benchmarked Prospector-Touchstone against other XL-MS search engines in the original study(45) including mass modification-based approaches (Prospector alone, XiSearch, and pLink2) and diagnostic-ion-centric methods (MeroX, MS-Annika, XlinkX, MaxLynx), by integrating Prospector-Touchstone analysis with the original results (Figure 2). The initial comparison was done searching a database of 171 *E. coli* proteins (ribosomal proteins and contaminants). When classifying URPs at the nominal 1% FDR reported by each pipeline, Prospector-Touchstone reports 654 URPs at a ground truth error of 2.7%. This is similar to the overall number reported by MeroX (658) although MeroX had a higher ground truth error rate (5.7%). MS-Annika and MaxLynx reported 632 and 635 URPs respectively with error rates of 2.7% and 2.2%. Hence, Prospector-Touchstone slightly outperformed the best of these selected pipelines on this classification task, demonstrating that mass modification algorithms can perform comparably to diagnostic-ion centric algorithms for DSSO analysis, without requiring detection of the signature ions. When reporting the search results at a corrected 1% ground truth FDR (Figure 2A-right panel), Prospector-Touchstone reports 609 URPs, 81 more than the next best strategy (MeroX).

**Figure 2.**
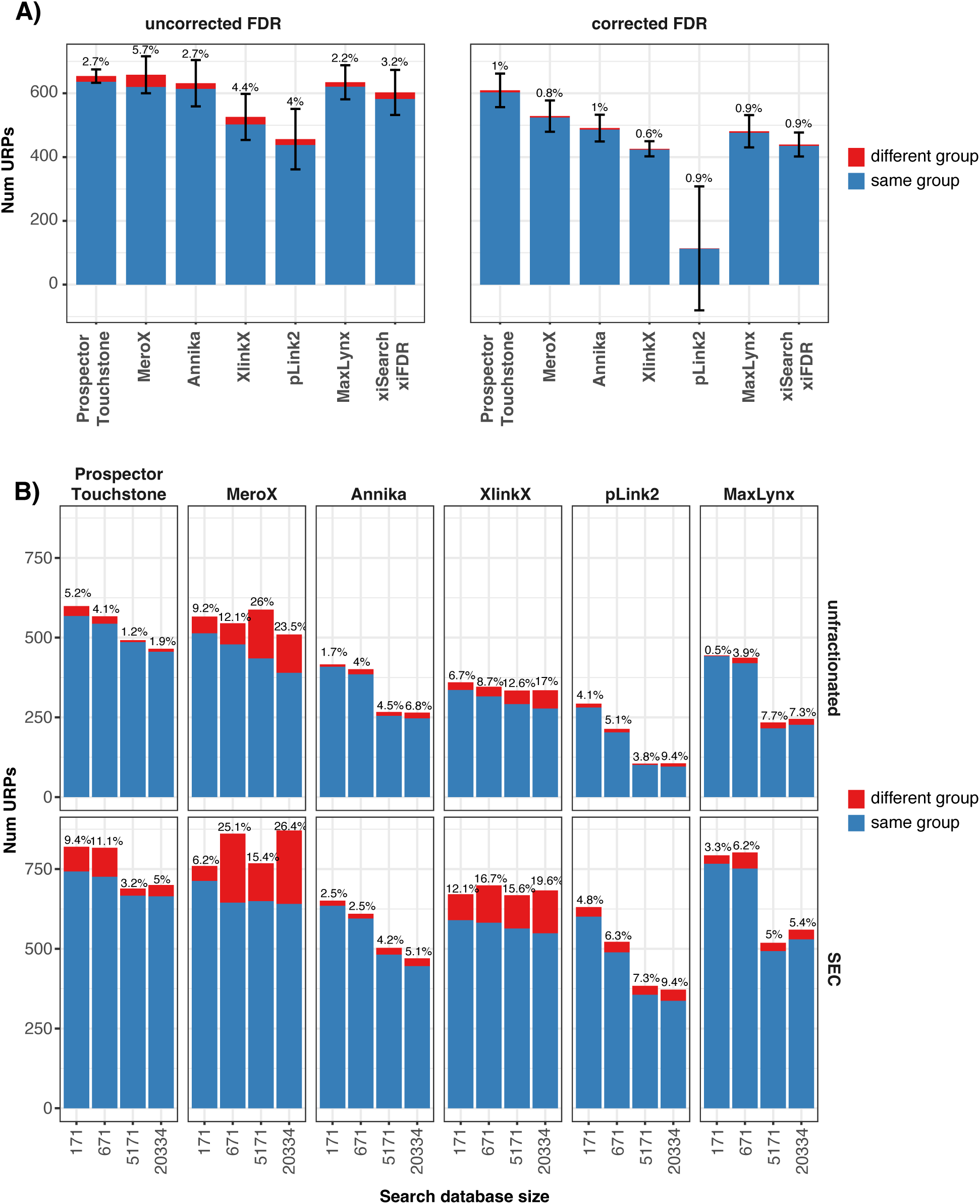
Software and Data availability. Benchmarking analysis against other mass modification based search engines (pLink2, xiSearch-xiFDR, Prospector-Touchstone) as well as algorithms which require diagnostic ions from cleavable reagents (MeroX, MS-Annika, XlinkX, MaxLynx) on a DSSO crosslinked synthetic ribosome library, acquired by MS2-stepped-HCD. All results are reported at a nominal 1% FDR calculated for unique residue pairs (URPs). The ground truth FDR was determined by tabulating number of disallowed crosslinks (between different experimental groups) relative to allowed crosslinks (within the same group) and reported above the bars on the plot. Experimental design was taken from Matzinger *et al.,* (2022) and the results using other bioinformatic platforms are taken directly from the source data of this reference. A) Average and standard deviation of three independent replicates of the main library. Data searched against 171 *E. coli* sequences comprising ribosomal subunits and abundant contaminants. Prospector – Touchstone analysis utilized a 10x scaled decoy database and a single threshold for all crosslinks. Left panel shows number of URPs reported at 1% nominal FDR (uncorrected). Right panel shows number of URPs reported at a 1% ground-truth FDR (corrected). B) Benchmarking against the main library spiked with 1:5 (by weight) linear HEK293 derived peptides. The search space included the 171 *E. coli* sequences with an additional 0, 500, 5000, or 20163, human sequences (number below bars). Prospector – Touchstone analysis utilized a 1x scaled decoy database and calculated separate thresholds for intra-protein and inter-protein crosslinks. The top panel shows results from a single unfractionated sample. The bottom panel shows results from 5 SEC fractions.

To mimic a more complex comparison, Matzinger *et al* used a library containing 1:5 spike-in of linear HEK293 derived peptides as background and performed database searches against the same 171 *E. coli* proteins as before plus 500, 5000, or 20163 human protein sequences respectively. Two versions of the dataset were searched to report results at 1% FDR at the URP level, one from the unfractionated and the other from size-exclusion chromatography (SEC) fractionated samples. To explore how database size affects error estimation and facilitate comparison, Prospector used a 1× decoy database for these larger searches, rather than the 10× decoy database used for the smaller Cas9 benchmark. Notably, FDR estimation accuracy improved as database size increased, 4.1-5.2% ground truth FDR when searching 171 or 671 sequences vs 1.2-1.9% when searching 5,171 and 20,334 sequences. This same trend is not apparent with the other search software. On the unfractionated data when searching with 20,334 sequences (Figure 2B-top), Prospector-Touchstone reported the highest number of correct URPs (456 correct and 9 incorrect) at the lowest error rate (1.9%) of all the search engines considered. MeroX which reported the highest number of total URPs did so at a 23.5% error rate (390 correct and 120 incorrect) and with fewer correct URPs than Prospector-Touchstone. These trends generally hold true across all the comparisons in Figure 2B including both the unfractionated and SEC fractionated data with Prospector-Touchstone reporting more URPs at lower error rates than comparative software. The only exception is that MaxLynx reports more URPs at lower error rates with the two smaller databases (171 and 671 proteins) in the SEC fractionated dataset, but not with the larger databases.

These results suggest Prospector-Touchstone often outperforms other leading platforms by identifying a greater number of crosslinks with high fidelity, particularly when searching large datasets comprising multiple LC-MS runs against entire proteomes. Note that, other than correctly calculating the theoretical product ions from cleavable crosslinkers, no other changes were made to score or otherwise consider the detection of diagnostic ion pairs. We also note that the version of pLink2 utilized in this comparison does not account for product ions from cleavable crosslinkers correctly and newer pLink versions that account for cleavable-linker product ions would be expected to improve performance in this analysis (49). As shown above, modifications to Prospector to correctly account for fragmentation of cleavable crosslinkers resulted in 1.5-2 fold more crosslinks detected in the DSSO data (Supplemental Figure 4).

### Assessing MS-acquisition strategies for optimal DSSO crosslink identification

With the Prospector-Touchstone platform established, we next sought to determine the optimal MS acquisition and data analysis strategies for identifying DSSO crosslinks in samples of different complexities. We evaluated nine MS acquisition strategies (methods m1-m9, Figure 3, Supplemental Table 1), including both commonly used single-stage fragmentation methods and workflows adapted from previously published XL-MS studies. These include single-stage electron-dissociation methods (m4: MS2.ETD and m5: MS2.EThcD) and collisional-dissociation methods (m6: MS2.HCD and m7: MS2.stHCD(34)), an MS3-based method (m2: MS2.CID-MS3.CID(30)), and hybrid workflows (m1: MS2.HCD-MS2.EThcD(3); m3: MS2.CID-MS3.CID-MS2.ETD(15); m8: MS2.CID-MS2.EThcD(31); and m9: MS2.CID-MS2.HCD(33)).

**Figure 3.**
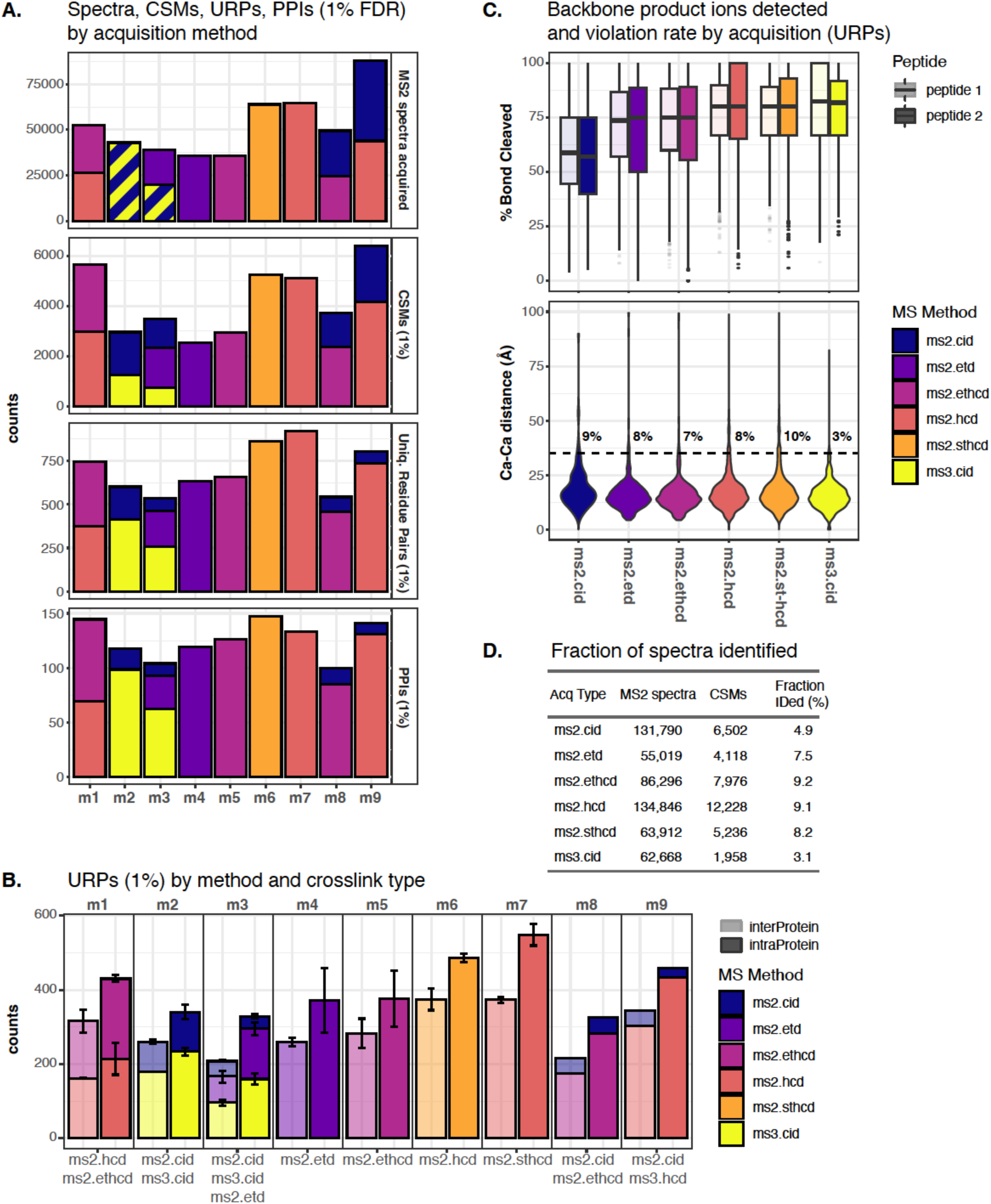
**Evaluation of MS acquisition strategies on DSSO crosslinked 80S ribosome**. Rabbit 80S ribosome was crosslinked with 0.5 mM DSSO and acquired by 9 different MS methods using an Orbitrap Lumos (m1-m9). A) Total product ion spectra acquired (MS2), CSMs classified at 1% FDR, and unique residue pairs (URPs) at 1% FDR, and protein-protein interactions (PPIs) at 1% FDR (note that intra-protein hits are omitted in the PPI analysis). The bars are colored by gas-phase acquisition method. CSMs are redundant because the same precursor was acquired by multiple methods in some cases. URPs are unique within the strategy and are assigned to the highest scoring CSM. B) URPs (1% FDR). Top panel shows the percentage of backbone product ions detected for the highest-scoring (peptide 1) and lowest-scoring (peptide 2) of the crosslinked peptide pair. Bottom panel, crosslinked spectral matches were mapped to a high-resolution ribosome structure (PDB ID = 6hcj) and Cα-Cα distances were measured. A distance greater than 35Å was considered a violation and the violation rate is reported above the bar. Note that many >35 Å crosslinks can be explained by ribosome aggregation (see text). C) URPs (1% FDR) reported by intra-protein and inter-protein pairs. D) The identification rate of different acquisition methods (percentage of potential product ion spectra successfully annotated as type 2 crosslinks (CSMs @ 1% FDR).

DSSO crosslinked 80S ribosomes were used to evaluate performance on a purified protein complex comprising 77 protein subunits (Supplemental Figure 5). MS2 data were searched using the Prospector-Touchstone pipeline while MS3 data were analyzed using the Prospector-XL-Tools(50) pipeline, which integrates MS3 search results with MS1 and MS2 data for high-confidence identification of DSSO crosslinks.

The highest number of MS2 spectra were obtained with sequential MS2.CID-MS2.HCD acquisition (m9) followed by MS2.HCD (m6) and stepped MS2.stHCD (m7) (Figure 3A). Notably, MS2.CID-MS2.HCD (m9) acquisition was much faster than other sequential methods such as MS2.HCD-MS2.EThcD (m1) and MS2.CID-MS2.EThcD (m8), due to the ability of the Lumos to parallelize the dissociation of isolated precursor ions, with CID taking place in the linear ion trap while HCD is carried out in the ion routing multipole. ETD-based acquisitions (m4 and m5) were considerably slower than HCD-based methods on the Lumos instrument used in this study, although this gap has closed considerably in newer generations of instruments (51). Methods that acquired MS3.CID data (m2, m3) were also relatively slow and acquired only slightly more MS2 spectra than the electron-transfer-based methods. For consistency, spectral acquisition and CSM counts for MS3-based methods were tabulated at the initiating MS2 level and an MS2 precursor event was considered successfully identified if it ultimately yielded a CSM, regardless of the number of contributing MS3 spectra.

In general, the number of crosslink assignments at different levels of summarization can be predicted by the number of distinct precursor ions acquired (Figure 3A). While m9 yielded more redundant CSMs, m6 and m7 characterized more unique precursors and consequently performed best among the acquisition strategies tested. At the URP level, m6 (861 ± 41 URPs) and m7 (922 ± 37 URPs) performed best, followed by the hybrid HCD-containing methods. MS3.CID and ETD-based workflows identified substantially fewer URPs (Figure 3A). The same pattern was observed at the protein-pair level. Across all three ETD/HCD comparisons (m4/m5 vs m6/m7, and m8 vs m9), replacing HCD with ETD reduced performance.

All nine methods identified more intra-protein than inter-protein residue pairs with intra-protein URPs exceeding inter-protein URPs by approximately 30-60%, as expected for the 80S ribosomal complex (22) (Figure 3B). The overall ranking of acquisition strategies remained largely unchanged when considering only inter-protein URPs. HCD-only methods (m6 and m7) identified the greatest number of inter-protein URPs, followed by the hybrid HCD-containing workflows (m9 and m1), whereas ETD-based and MS3-based methods produced substantially fewer identifications (Figure 3B).

For the two workflows incorporating MS3 analysis (m2 and m3), a substantial fraction of accepted inter-protein URPs originated from the accompanying MS2 acquisitions rather than the MS3 spectra themselves. For m2, 180 ± 1 inter-protein URPs were identified exclusively from the MS3.CID analysis, whereas an additional 81 ± 6 were contributed by the MS2.CID spectra. Similarly, for m3, the MS3.CID analysis contributed 96 ± 7 URPs, while the accompanying MS2.CID and MS2.ETD acquisitions identified an additional 40 ± 2 and 73 ± 16 URPs, respectively. These results demonstrate that searching the accompanying MS2 acquisitions could be beneficial to increase the overall cross-link identification of MS3 workflows in a purified protein complex.

### Fragmentation Characteristics Underlying Acquisition Performance

To understand the differences in performance among the acquisition strategies, we compared the fragmentation characteristics of the individual dissociation methods by quantifying the fraction of peptide bonds cleaved in each of the two constituent peptides of the crosslink (Figure 3C). As expected, MS2.CID spectra, which were acquired under conditions optimized to generate DSSO diagnostic ions rather than backbone fragmentation, exhibited the lowest backbone cleavage efficiency of all the dissociation types. More surprisingly, ETD-based methods also generated fewer backbone cleavages from DSSO-crosslinked peptides than HCD-based methods, despite their superior performance with non-cleavable crosslinkers. In a previously published BS3 dataset of similar complexity, electron dissociation produced both a higher fragmentation yield and more symmetric fragmentation of the two constituent peptides than HCD (3) (Supplemental Figure 6).

This observation is consistent with the findings of Rappsilber and colleagues(52) that HCD enhances backbone fragmentation of cleavable linkers relative to non-cleavable linkers by improving the yield of product ions that contain the crosslinked amino acid residue. In non-cleavable crosslinkers these ions retain the mass of the second peptide, making them more difficult to detect by HCD and therefore favoring ETD. Because electron-transfer does not cleave DSSO, it provides no analogous advantage for MS-cleavable crosslinkers. EThcD produced similar median fragmentation efficiencies for DSSO (75% / 75%) and BS3 (80% / 75%) for peptide.1/peptide.2 (Figure 3C, Supplemental Figure 6). In contrast, HCD fragmented substantially more efficiently with DSSO (80% / 80%) than BS3 (75% / 57%) from both peptides. Overall, the enhanced HCD fragmentation afforded by DSSO largely eliminates the historical advantage of electron-transfer dissociation for confident crosslink identification. The improvement is particularly evident for peptide.2, the less extensively fragmented peptide, which is often the limiting factor in successful crosslink assignment (35).

Although MS3.CID generated the greatest backbone fragmentation efficiency, only 3.1% of acquired MS2 precursor events ultimately yielded accepted CSMs, compared with 9.1% for MS2.HCD acquisitions (Figure 3D). The lower identification rate of MS3-based workflows is therefore not attributable to fragmentation quality, but rather to their longer cycle times, and the need to accurately select diagnostic ions for subsequent MS3 analysis. In addition, cross-link identification requires stringent integration of MS1, MS2, and MS3 information, including correct identification of both cross-linked peptide constituents. Together, these constraints reduce CSM recovery while providing greater confidence in the resulting identifications. In contrast, HCD-only MS2 workflows combine rapid spectral acquisition with a high identification rate, resulting in greater recovery of biologically informative URPs and protein-pairs, albeit with a higher error rate.

### Structural Validation of Acquisition Strategies

We next compared the consistency of the different acquisition strategies with the high-resolution structure of the rabbit 80S ribosome (PDB: 6HCJ) (Figure 3C, bottom). Most mapped URPs from every method corresponded to Cα–Cα distances below 35 Å. However, the fraction of apparent distance violations varied from 2.6% for MS3.CID to 9.7% for MS2.stHCD. Notably, many of the long-distance crosslinks identified by MS2.stHCD were high-quality matches involving residues separated by more than 90 Å in the monomeric ribosome structure (Supplemental Figure 7). Most of these apparent violations occur between residues on the 40S subunit and are consistent with previously described collided ribosome dimers(53) (Supplemental Figure 8). Low levels of ribosome aggregation were also observed by negative-stain EM under the DSSO crosslinking conditions (Supplemental Figure 5A). We therefore attribute the higher violation rate of HCD methods primarily to their greater sampling depth, which enables detection of crosslinks arising from conformationally heterogeneous or low-abundance species, rather than to reduced identification accuracy. This interpretation is supported by three observations: (i) the scoring and FDR strategy was extensively validated using the benchmark datasets (Figures 1 and 2); (ii) HCD-based methods generated the greatest backbone fragmentation among the MS2 acquisition methods (Figure 3C); and (iii) manual inspection confirmed that many long-distance crosslinks were supported by high-quality spectra (Supplemental Figure 7). In contrast, MS3-based identifications must satisfy additional criteria relative to MS2-based classifications and typically exhibit empirical decoy rates well below the nominal 1% acceptance threshold. The low structural-violation rate observed for MS3.CID therefore likely reflects both reduced sampling of low-abundance conformations and a lower classification error rate.

### Diagnostic-ion filtering does not improve crosslink recovery

The Prospector-Touchstone scoring and classification workflow does not explicitly require the diagnostic ion pairs produced by MS-cleavable crosslinkers such as DSSO. We therefore tested whether diagnostic-ion detection could improve DSSO crosslink identification using two distinct approaches: prefiltering spectra before database searching and incorporating diagnostic-pair detection into post-search re-scoring and classification.

For the first approach, MS-Filter, a Prospector utility for retaining spectra that contain certain specified fragment masses or mass relationships(54), was extended to detect product-ion pairs corresponding to the expected DSSO cleavage products. A DSSO-crosslinked precursor can produce alkene-and thiol-modified products from both constituent peptides. MS-Filter therefore detects pairs of ions whose masses sum to the precursor mass with the expected offsets for thiol/thiol, alkene/thiol, or alkene/alkene products (Supplemental Figure 9).

DSSO diagnostic ions formed most efficiently under CID and were produced over a broader collision-energy range than with HCD (Supplemental Figure 10). They are not formed efficiently by electron transfer. For hybrid workflows, diagnostic-pair status was determined from the linked MS2.CID spectrum and assigned to the corresponding HCD, ETD, or EThcD spectrum acquired from the same precursor. For single-stage dissociation workflows, diagnostic-ion filtering was applied directly to the acquired spectrum.

We compared searches of filtered and unfiltered DSSO-crosslinked ribosome datasets (Figure 4A). Diagnostic-ion filtering reduced the numbers of classified CSMs, URPs, and protein pairs for nearly every acquisition strategy. For instance, for

**Figure 4.**
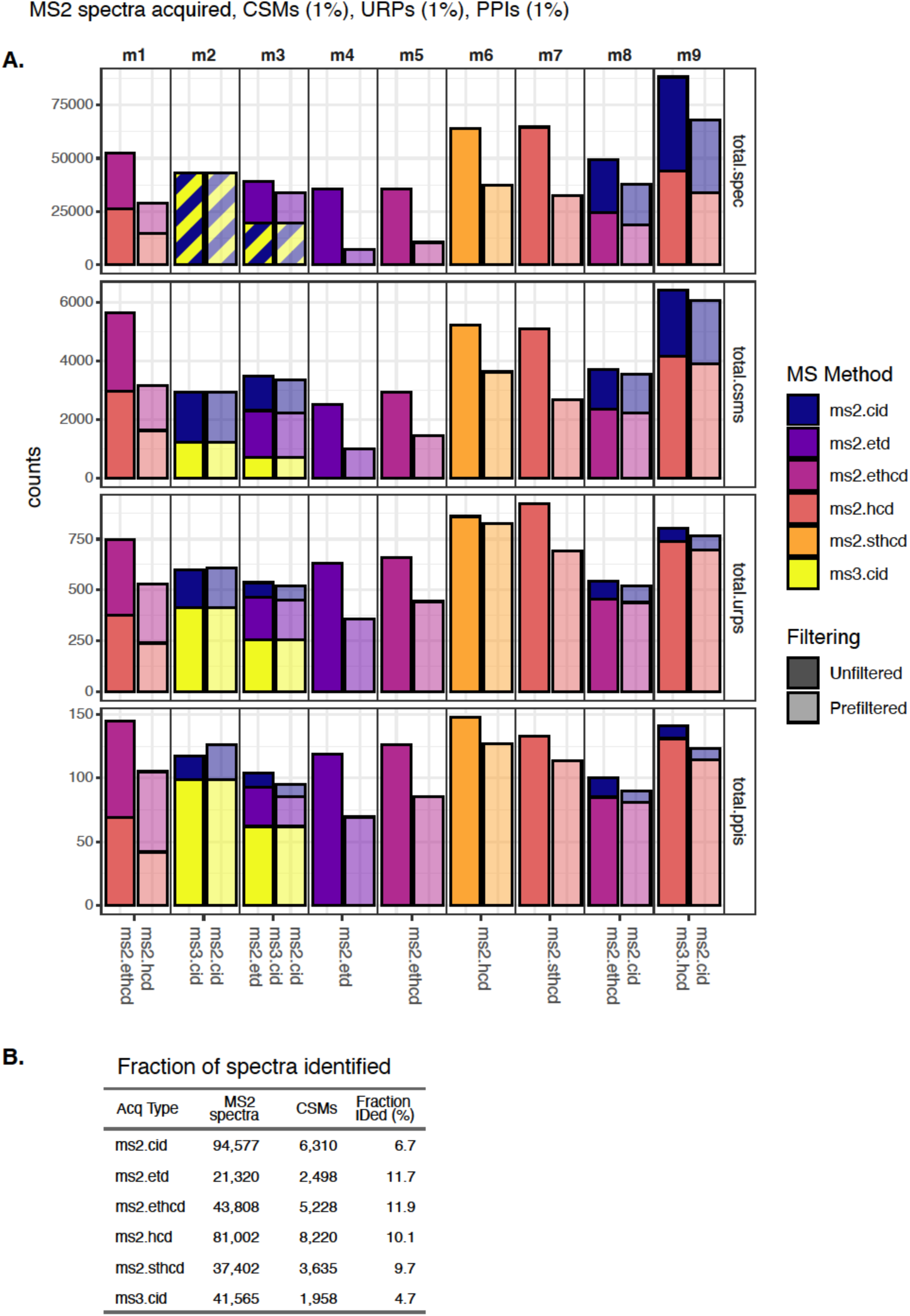
Analysis of DSSO crosslinked 80S ribosome using diagnostic ion filtering shows decreased performance. MS-Filter was used to create filtered peaklists based on detection of diagnostic ion pairs in the MS2 analysis. For solo methods, the diagnostic ions were detected in that scan type. For hybrid methods with an MS2-CID scan, linked MS2-HCD or MS2-ETD scans were retained if the diagnostic ion pairs were detected in the MS2-CID scan. Results were reported as in Figure 3A.

MS2.stHCD, filtering reduced the number of protein pairs from 148 ± 2 to 127 ± 1. Reductions were observed for all HCD-based workflows including hybrid methods in which diagnostic-pair detection was obtained from the linked CID scan. As expected, the largest losses occurred for ETD-based methods, which inefficiently cleave the DSSO sulfoxide bond. The only exception was MS2.CID-MS3.CID, which showed a modest increase in protein-pair identifications after filtering, with the increase arising entirely from the MS2.CID portion of the analysis. Thus, requiring detectable diagnostic-ion pairs generally reduced rather than improved crosslink recovery.

We also calculated the fraction of searched spectra that were identified as CSMs (Figure 4B, compare with Figure 3D). Diagnostic-ion filtering generally increased the identification rate among retained spectra, indicating that detectable diagnostic pairs enrich for spectra derived from identifiable crosslinked peptides. However, this enrichment was insufficient to compensate for the reduction in spectra submitted for searching, and fewer CSMs were identified overall. For example, Prospector-Touchstone classified 6,502 CSMs from 131,790 unfiltered MS2.CID spectra, corresponding to an identification rate of 4.9%. Filtering retained 94,577 spectra and yielded 6,310 CSMs, increasing the identification rate to 6.7% while reducing the total number of identifications. Thus, even under MS2.CID conditions optimized for DSSO cleavage, requiring detectable diagnostic-ion pairs excludes a subset of spectra that can otherwise support confident CSM assignments.

### Diagnostic ions do not improve discrimination of correct and incorrect CSMs

To determine why diagnostic-ion filtering reduced performance, we examined diagnostic-pair detection among target and decoy CSMs above and below the classification threshold (Figure 5, Supplemental Figure 11, Supplemental Tables 2 and 3). A spectrum was considered diagnostic-pair positive when it contained an alkene/alkene, alkene/thiol, or thiol/thiol pair detected across the permitted charge states and exceeding a minimum abundance threshold of 5% of total matched ion intensity. Lower-abundance diagnostic ions may therefore have remained undetected.

**Figure 5.**
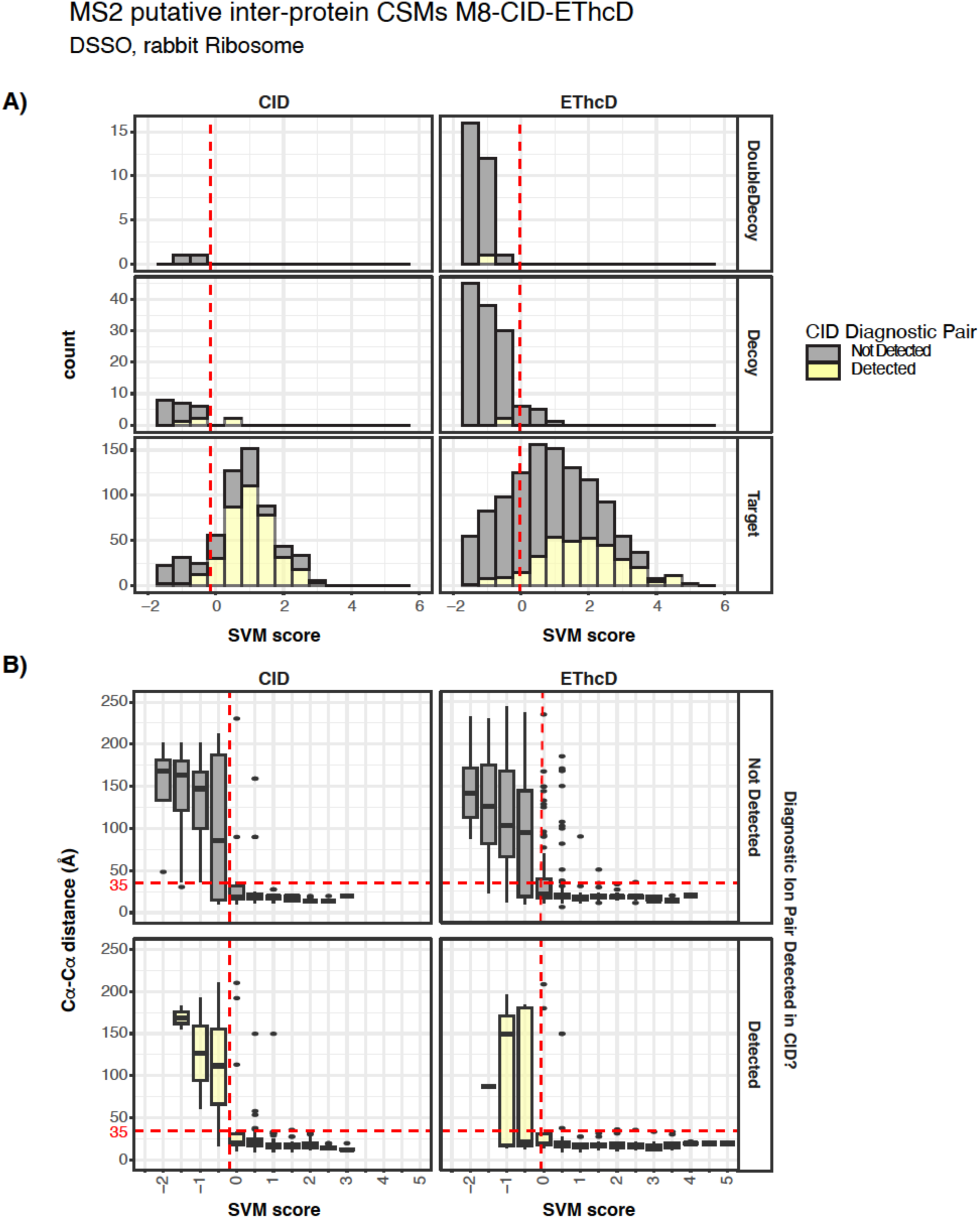
Distribution of diagnostic ion pairs in linked CID and EThcD spectra from 80S ribosome. A) Diagnostic alkene-alkene, alkene-thiol, or thiol-thiol ion pairs from DSSO crosslinking are distributed similarly in putative inter-protein hits from target and decoy databases. The red dashed line shows the acceptance SVM score threshold for 1% FDR CSM classification. Left panels show CID data with diagnostic pairs detected in the CID scan. Right panels show EThcD data with diagnostic pairs detected in the linked CID scan. Yellow indicates that a diagnostic ion pair was detected in the hit, while grey bars indicate that the pair was not detected. B) The diagnostic ion pairs are distributed similarly in correct and incorrect target database inter-protein putative hits as measured by mapped Cα distances on the rabbit 80S ribosome high resolution structure (PDBid = 6hcj). Vertical dashed red lines show the 1% FDR CSM classification threshold. Horizontal lines show 35Å Cα-Cα threshold used to define a violation.

Diagnostic pairs were present in approximately 72–74% of accepted target MS2.CID CSMs, but also in approximately 78–82% of decoy matches above the classification threshold (Figure 5, Supplemental Figure 11, Supplemental Tables 2 and 3). Among lower-scoring matches, diagnostic pairs were modestly enriched in target relative to decoy hits, but their presence did not cleanly separate correct from incorrect assignments. Consequently, filtering rejects some correct CSMs lacking detectable diagnostic ions while retaining incorrect CSMs that contain them.

We next asked whether diagnostic-ion detection could identify a population of correct target CSMs immediately below the 1% FDR threshold. Cα-Cα distance distributions showed that the scoring threshold already separated most structurally consistent and inconsistent assignments without considering diagnostic ions (Figure 5B; Supplemental Figure 11B, left panels). Among CSMs immediately below the threshold, distance distributions were similar for spectra with detected diagnostic pairs (yellow bars) and without detected diagnostic pairs (grey bars). Diagnostic-ion detection therefore did not identify a distinct population of structurally supported CSMs below the default score threshold that could be rescued by incorporating diagnostic-pair evidence.

The same behavior was observed for multi-stage activation strategies in which diagnostic-pair status was measured in a linked MS2.CID scan and peptide assignment was supported by the corresponding HCD or EThcD spectrum (Figure 5B; Supplemental Figure 11B, right panels). In these datasets, diagnostic pairs were detected in the CID scans of both structurally consistent and distance-violating assignments, while some structurally consistent assignments lacked detectable diagnostic pairs. Nevertheless, the Touchstone scoring model effectively discriminated correct from incorrect assignments using peptide fragment ions detected in the HCD or EThcD spectra, regardless of diagnostic pair detection.

Incorporating diagnostic-pair detection directly into the scoring function also failed to improve classification. Scores that either added an arbitrary diagnostic-ion contribution (“diagScore.1-3”) or included diagnostic-pair detection as an SVM feature (“SVM.diagnostic”) classified fewer CSMs than the default Touchstone score (“SVM.score”) throughout the low-FDR range of interest (Supplemental Figure 12). Prospector does assign conventional peptide-spectrum score to diagnostic ions, as it does to P, PL, and PLK ions observed from nominally non-cleavable reagents such as BS3 and DSS (35). Diagnostic-ion matches can therefore affect features such as “Score Difference” and the “% of TIC matched”. However, Prospector does not explicitly reward the occurrence of a complementary diagnostic-ion pair.

The limited discriminatory value of diagnostic-ion pairs reflects the nature of the crosslink search problem. For most ambiguous CSMs, the better-fragmented peptide is assigned correctly and uncertainty lies in the identity of the more weakly fragmented peptide (35). When several candidate crosslinks share the same assignment for peptide.1 and precursor mass, mass balance dictates the same complementary diagnostic-ion masses. Thus, diagnostic ions can confirm the presence of a cleaved crosslink precursor but cannot distinguish among competing assignments for peptide.2, which instead requires sufficient sequence-specific product ions. Supplemental Figure 13 illustrates this problem using an MS2.CID spectrum from the ribosome dataset.

Three candidate assignments share the same peptide.1 sequence from eL28 but assign peptide.2 to different intra-or inter-protein partners. All three candidates match the precursor mass and the four expected alkene-and thiol-modified diagnostic ions. The correct assignment is distinguished only by a sequence-specific backbone fragment from peptide.2.

Thus, even when diagnostic-ion pairs are explicitly incorporated into the score, sequence-specific product ions are more informative for distinguishing correct from incorrect CSMs. Diagnostic ions provide evidence that a cleavable crosslinked precursor was fragmented, but they do not necessarily identify the two constituent peptides. Requiring their detection can also exclude valid spectra in which diagnostic ions are weak or absent. Because diagnostic-centered search strategies require diagnostic-ion matches by design, they cannot report the valid assignments lost through this requirement.

### Performance of different acquisition strategies in proteome-wide XL-MS

To evaluate whether the trends observed with the purified ribosome extended to proteome-scale XL-MS, we analyzed DSSO-crosslinked HEK293 whole-cell lysates (WCL) using three acquisition strategies: MS2.EThcD (m8), MS2.stHCD (m7), and MS2.CID-MS3.CID (m2). The MS2 datasets were analyzed using Prospector-Touchstone, whereas the MS3 workflow was analyzed using Prospector-XL-Tools, with crosslinks identified from the accompanying MS2.CID spectra combined with those reconstructed from the MS3.CID data.

Consistent with the ribosome experiments, MS2.stHCD acquired the greatest number of spectra and produced the largest number of CSMs (Figure 6A). MS2.stHCD identified 46,411 intra-protein and 3,373 inter-protein CSMs at 1% FDR, compared with 27,078 intra-protein and 2,206 inter-protein CSMs from MS2.CID-MS3.CID. From the latter category, 1,655 inter-protein CSMs were classified from the MS3.CID and 551 CSMs were added by considering the MS2.CID scans. MS2.EThcD performed between these two methods at the spectral level with 31,461 intra-protein and 2,525 inter-protein CSMs.

**Figure 6.**
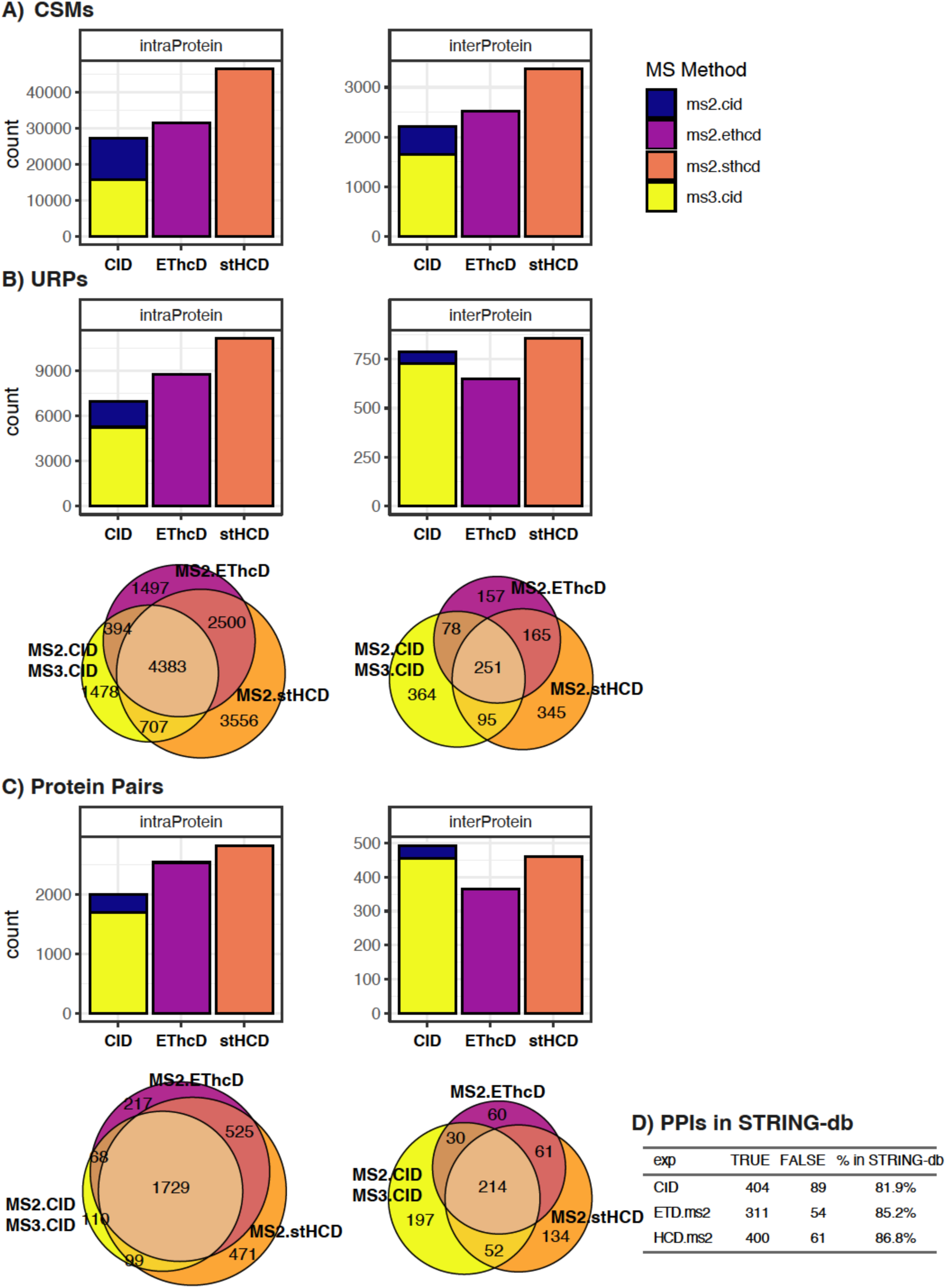
Evaluation of MS-acquisition strategies on DSSO crosslinked HEK293 cell lysates. HEK293 whole cell lysates were crosslinked with DSSO and analyzed using three representative MS methods. Data were classified at 1% FDR using separate thresholds for inter-protein and intra-protein crosslinks using Prospector-Touchstone (for MS2 level analysis) or Prospector-XLtools (for MS3 level analysis). Intra-protein hits and MS3-level results have actual FDRs much lower than the 1% nominal classification threshold. Results are shown at the A) CSM, B) URP, and C) PP level, with Venn Diagrams showing intersections of the various methods for URPs and PPs. D) Percentage of PPIs (inter-protein PPs) in which the two linked proteins are annotated in STRING-db with score > 900.

The same overall trend was observed at the residue-pair level. MS2.stHCD classified 11,146 intra-protein and 856 inter-protein URPs, compared with 6,962 and 788, respectively, from the MS2.CID-MS3.CID workflow (Figure 6B). Of the 788 inter-protein URPs, 728 were identified by MS3.CID, with only 60 additional URPs contributed by the accompanying MS2.CID scans. At the protein-pair level, however, MS2.CID-MS3.CID reported 493 inter-protein protein pairs, approximately 5% more than the 461 identified by MS2.stHCD (Figure 6C). Thus, while MS2.stHCD provided greater overall CSM and URP coverage, MS2.CID-MS3.CID yielded slightly more inter-protein PPIs. Notably, MS3.CID alone identified most of these interactions (455 of the 493 PPIs), with the accompanying MS2.CID analysis adding only 38 protein pairs. Collectively, these results suggest that MS2.stHCD and MS3.CID workflows are complementary in proteome-wide studies.

### Error characteristics and biological support of MS2-and MS3-based results

We next compared error propagation in the MS2-and MS3-based analyses. In the MS3-centric analysis, linear MS3 peptide assignments are first classified using a conventional target-decoy search and are then integrated with the corresponding MS1 and MS2 information to reconstruct crosslink matches. For a linear MS3 decoy assignment to propagate into a decoy crosslink match, the peptide must contain a DSSO fragment on an internal lysine, a linked MS3 spectrum must identify a second peptide with a DSSO fragment, and the two peptides together must be consistent with the MS1 precursor ion. Most decoy hits from the initial MS3 search failed these criteria. In the HEK293 dataset, no decoy MS3 assignments propagated to the CSM, URP, or protein-pair levels, even when the initial MS3 classification criteria were broadened to generate additional linear decoys.

MS2-based mass-modification searches produced substantially more decoy candidates. These searches initially identify linear peptides bearing an arbitrary mass modification on an internal lysine. Because the mass modification is scanned through a broad window (typically 500–4,000 Da in discrete steps), hundreds to thousands of candidate assignments may be retained for each peptide. Crosslink reconstruction then evaluates combinations of candidate peptide assignments whose mass modifications are mutually consistent with the mass of the other peptide and the DSSO bridge. By contrast, MS3 reconstruction typically considers only the small number of peptides independently identified from linked MS3 spectra.

Because Touchstone classifies inter-and intra-protein assignments separately and the intra-protein MS2 searches produced too few decoy assignments for stable estimation of a 1% FDR threshold, the reported 1% FDR thresholds for the MS2 datasets refer specifically to inter-protein CSMs, URPs, and protein pairs. No decoy MS3 assignments were observed at these levels in the HEK293 dataset, corresponding to a nominal empirical decoy rate of 0%.

To determine whether differences in yield and error control affected biological validity, we compared the identified inter-protein protein pairs with STRING-db annotations(55). The fraction supported by high-confidence STRING interactions (score > 0.9) was similar for all three methods: 81.9% for MS2.CID-MS3.CID, 85.2% for MS2.EThcD, and 86.8% for MS2.stHCD (Figure 6D). Thus, despite differences in acquisition speed, identification yield, and empirical error behavior, all three workflows predominantly recovered well-supported protein interactions.

The three methods also showed substantial overlap at the intra-protein protein level. Fifty-four percent of intra-protein protein-level identifications were detected by all three methods, and only a small fraction of the proteins identified by MS2.CID-MS3.CID or MS2.EThcD were absent from the larger MS2.stHCD dataset (Figure 6C). In contrast, only 29% of inter-protein protein pairs were shared across all three datasets. This low overlap is consistent with the greater stochasticity of sampling lower-abundance inter-protein crosslinks and with differences in fragmentation and identification efficiency among the acquisition strategies. The relative proportions of intra-and inter-protein identifications also differed among the workflows. Inter-protein protein pairs accounted for 20.6% of protein-level identifications from MS2.CID-MS3.CID, compared with 14.4% from MS2.stHCD.

### MS data acquisition and cross-link identification efficiency

To distinguish acquisition speed from identification efficiency, we examined the fate of spectra acquired by the MS2.stHCD and MS2.CID-MS3.CID workflows (Figure 7).

**Figure 7.**
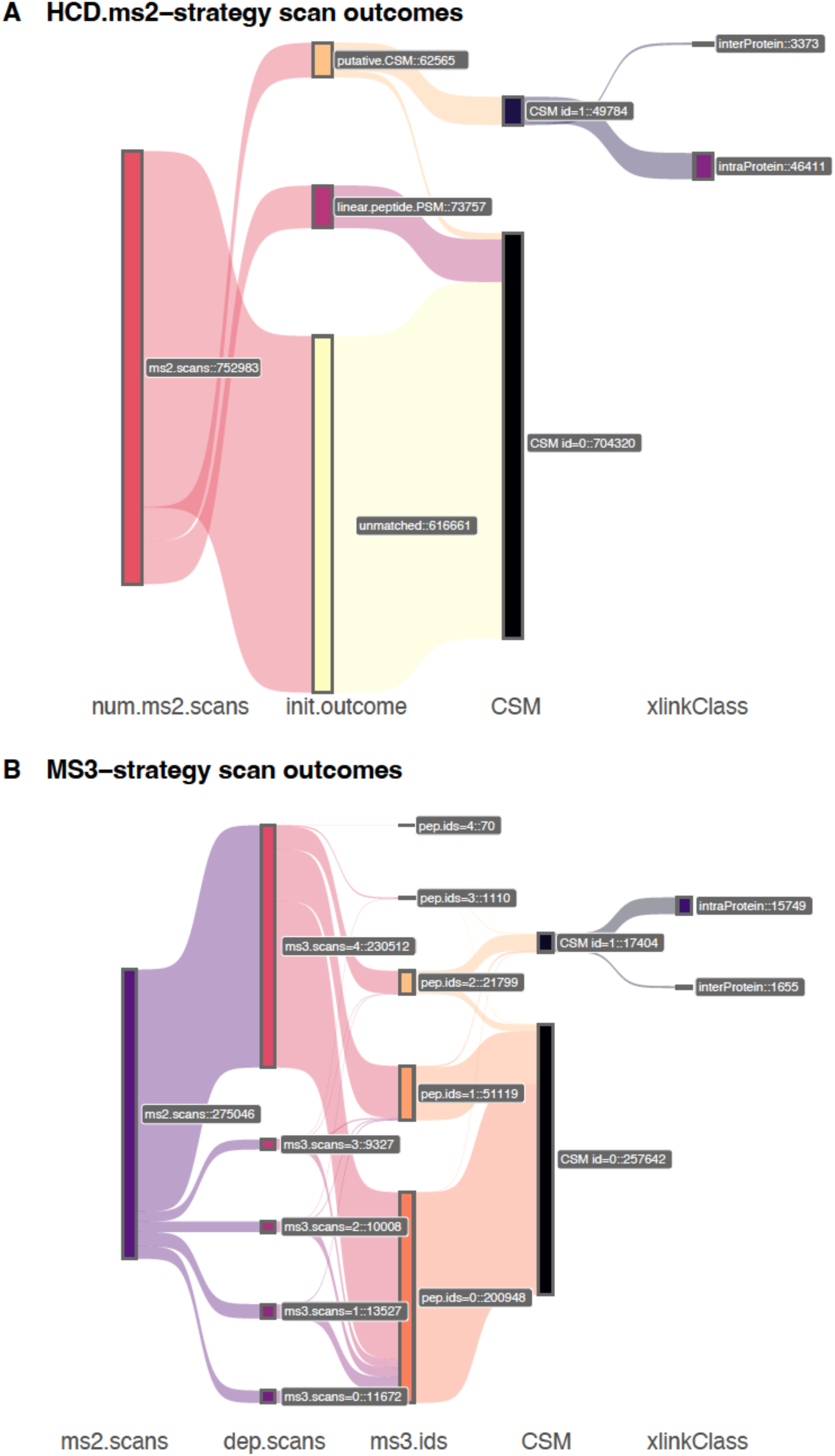
Outcomes of all MS2 spectra acquired on HEK293 cells from MS2.stHCD and MS3-CID analysis. Sankey flow diagrams tracing the fate of A) 752,983 MS2.stHCD scans and B) 275,046 MS2-CID scans acquired on the same DSSO crosslinked HEK293 WCL sample analyzed by either Prospector-Touchstone or Prospector-XLtools respectively. The Sankey diagrams trace how many spectra are initially identified as linear peptides or putative crosslinked peptides before being validated as authentic CSMs by Touchstone. For MS3, the diagram traces how many MS3 scans are triggered by each MS2 scan, whether those MS3 scans are matched to linear peptide fragments, and whether those peptide matches can be reconstituted into an authentic CSM.

Across all fractions and replicates, MS2.stHCD acquired 752,983 MS2 spectra. Of these, 62,565 were assigned as putative crosslinks and 73,757 were matched to linear peptides, while 82% remained unidentified. Following Touchstone classification, 49,784 putative crosslinks were accepted as CSMs at 1% FDR, including 46,411 intra-protein and 3,373 inter-protein CSMs. Relative to all acquired spectra, this corresponded to identification rates of 6.2% for intra-protein and 0.4% for inter-protein CSMs.

In contrast, MS3.CID analysis required additional stages of precursor selection, fragmentation, and ion detection. Consequently, the workflow acquired only 275,046 initiating MS2 spectra, approximately threefold fewer than MS2.stHCD. Each MS2 spectrum triggered between zero and four dependent MS3 scans, with 83.8% triggering the maximum of four. Among the initiating MS2 precursor events, 21,799 yielded identifications of at least two peptides from the associated MS3 spectra and could therefore potentially be reconstructed as crosslinks. An additional 51,119 events yielded only one identified peptide, suggesting that many were crosslinked precursors for which the second constituent peptide could not be identified. Of the events yielding two or more peptide-spectrum matches, 75.7% were reconstructed as CSMs, resulting in 15,749 intra-protein and 1,655 inter-protein CSMs. These corresponded to overall identification rates of 5.7% and 0.6%, respectively, relative to the number of initiating MS2 precursor events.

Thus, the two workflows had broadly similar per-precursor identification efficiencies. MS3 analysis produced a slightly higher rate of inter-protein CSM identification and a somewhat lower rate of intra-protein identification. These comparatively small differences were outweighed by two larger factors. First, the additional MS3 acquisition stages reduced the number of precursor ions sampled by approximately threefold.

Second, MS3-based identifications exhibited a substantially lower empirical decoy rate than the MS2-based results. The choice between the methods therefore reflects a tradeoff between the greater sampling depth of MS2.stHCD and the more restrictive, lower-error identifications produced by MS3 analysis.

An acquisition strategy that selectively triggered MS3 only for spectra likely to benefit from additional fragmentation could potentially retain the error-control advantages of MS3 while reducing its acquisition-time penalty. This could be implemented via real time search-based logic (56, 57) or by detecting features in the MS2 or MS1 scans that trigger additional MS3 scans. We therefore examined whether precursor or peptide characteristics distinguished CSMs identified by the three acquisition methods (Supplemental Figures 14 and 15). Peptide-length and precursor-*m/z* distributions were broadly similar among the datasets. Consistent with the preference of electron-transfer dissociation for precursors with higher charge density, MS2.EThcD identified proportionally more 5+ precursors and performed less well for 4+ precursors above 700 *m/z*. MS2.CID-MS3.CID identified proportionally fewer precursors with charge states of 5+ or higher and more 4+ precursors than the other methods. Nevertheless, MS2.stHCD identified the greatest number of CSMs across all charge states examined, and these basic precursor characteristics did not reveal a population that could be readily selected for preferential MS3 acquisition.

## Discussion

### Prospector–Touchstone enables accurate MS2-based XL-MS analysis

Recent community-wide benchmarking studies have highlighted substantial discrepancies between reported and true false discovery rates in XL-MS analyses, particularly at downstream levels of data summarization, underscoring the need for robust error-estimation and classification strategies (58). Here, we developed Prospector-Touchstone as an integrated pipeline for MS2-based identification of both cleavable and non-cleavable crosslinks with explicit FDR control at the CSM, unique residue-pair, and protein-pair levels. Protein Prospector was among the first generation of XL-MS search engines to employ a mass modification algorithm to linearize the search for crosslinked peptides (35, 41). To support MS-cleavable reagents, we introduced a cleavable-crosslinker class that calculates backbone product ions containing the defined remnants generated by cleavage of DSSO and similar reagents. Touchstone then performs post-search scoring, classification, summarization, and error estimation. Together, these developments provide a general platform applicable to samples ranging from purified complexes to proteome-scale preparations.

Leveraging two previously published XL-MS benchmark studies (44, 45), we have demonstrated that Prospector-Touchstone combines high identification yield with well-calibrated FDR control. For the non-cleavable BS3 benchmark, it classified more CSMs than other pipelines, in part because its reported error rate more closely approached the intended threshold. When URP yields were compared at a corrected 1% FDR to account for differences in calibration among software tools, Prospector-Touchstone reported the second-highest number of URPs. For the cleavable DSSO benchmark, it reported the largest number of URPs at a corrected 1% FDR and provided the most accurate residue-pair-level FDR estimates for the more complex searches containing HEK293 lysate background. Notably, Prospector-Touchstone also performed well in a recent inter-laboratory comparison organized by the Rappsilber group, in which software developers analyzed an extensively fractionated DSSO-crosslinked *E. coli* dataset and reported results at a nominal 2% FDR (58). The experimental design incorporated both an entrapment database to estimate the true error rate and protein-correlation profiling to provide an orthogonal assessment of protein interactions.

Prospector-Touchstone showed strong agreement between its reported FDR and these independent estimates. Taken together, these benchmark studies demonstrate that the Touchstone pipeline provides robust error control across different crosslinking chemistries, database sizes, and sample complexities. The established platform represents a substantial improvement over the earlier stand-alone Prospector workflow and performs comparably to or better than the top-performing software evaluated in the published studies (44, 45). Its ability to analyze both cleavable and non-cleavable cross-link data broadens the utility of Prospector-Touchstone across diverse XL-MS applications.

### Mass-modification searching remains effective for cleavable crosslinkers

Crosslink search engines can be broadly divided into approaches that require crosslinker-derived diagnostic ions to determine the constituent peptide masses and open-search approaches that identify candidate peptides without initially requiring their masses to account for the intact precursor. Diagnostic-ion-centric searches reduce the candidate space and can therefore be computationally efficient, but they generally require detectable and correctly assigned signature ions. Mass-modification searches do not depend on diagnostic-ion detection and can recover crosslinks from spectra in which these ions are weak, absent, or incorrectly annotated, although this broader search requires substantially greater computation and more extensive post-search classification.

Mass-modification algorithms were developed largely before the widespread adoption of MS-cleavable reagents, and their suitability for proteome-scale cleavable-linker analysis has therefore been less thoroughly explored. The performance of Prospector-Touchstone in the DSSO benchmarks demonstrates that mass-modification searching remains well suited to cleavable-linker analysis and can perform competitively even against algorithms designed specifically around diagnostic-ion detection. Recent proteome-scale XL-MS studies from the Rappsilber and Urlaub groups have likewise used mass-modification-based search engines (17, 42).

The principal disadvantage of open search strategies is computational cost. In the present study, the most intensive proteome-wide searches required either a 64-core server or up to 256 slots in a high-performance computing environment and typically completed within three to four days. We have also deployed Prospector successfully in a commercial cloud-computing environment. Although these requirements are substantially greater than those of MS3-based linear-peptide searches, the necessary computing resources are increasingly accessible, and further parallelization and optimization should continue to expand the practical scope of mass-modification analysis.

### Diagnostic ions identify cleavable crosslinked precursors but do not improve assignment specificity

We tested several approaches for leveraging DSSO diagnostic-ion pairs in Prospector-Touchstone analysis, including prefiltering spectra and incorporating diagnostic-pair detection into post-search scoring. Diagnostic-ion filtering enriched the retained peak lists for spectra that yielded accepted CSMs, but this enrichment was insufficient to compensate for the valid spectra that were discarded, leading to lower overall recovery. Similarly, adding diagnostic-pair information to the Touchstone scoring function reduced rather than improved classification performance. Although DSSO diagnostic pairs were detected in approximately 70% of accepted inter-protein MS2.CID CSMs, they were also common in decoy assignments and in over-length target matches. Their presence therefore cannot cleanly distinguish correct from incorrect peptide-pair assignments. This result can be rationalized by the fact that diagnostic ions primarily indicate cleavage of an MS-cleavable cross-linked precursor, whereas correct cross-link identification relies on sequence identification of the two peptide constituents.

Insufficient backbone fragmentation can therefore lead to incorrect CSM assignments despite the presence of diagnostic ions. Conversely, requiring diagnostic-ion detection can exclude otherwise valid MS2 spectra in which diagnostic ions are weak or undetectable.

This helps explain why Prospector-Touchstone performed favorably in the DSSO benchmark relative to diagnostic-ion centered platforms. It also highlights an inherent difficulty in evaluating diagnostic-ion-centric search strategies: because diagnostic-ion evidence is required before a putative match is considered, their reported results cannot reveal the valid assignments excluded by that requirement.

### DSSO changes the relative advantages of HCD and electron-transfer dissociation

An important benefit of MS-cleavable reagents is enhanced backbone fragmentation rather than diagnostic-ion production alone. With non-cleavable reagents, electron-transfer-based dissociation, particularly EThcD, can provide more extensive and balanced fragmentation of the two linked peptides than HCD for charge-dense precursor ions (59). Large product ions containing the intact crosslink and the mass of the second peptide are rarely detected after HCD, likely because they undergo further dissociation or are detected inefficiently, whereas electron-transfer dissociation can preserve and detect these large products more effectively. This has historically favored ETD or EThcD for non-cleavable XL-MS, where the richer fragmentation could compensate for slower acquisition rate.

DSSO changes this balance because collisional activation cleaves the sulfoxide bond, allowing backbone ions to retain only a small residual linker mass. HCD therefore provides strong sequence coverage from both linked peptides. The improvement is particularly important for peptide.2, which is less extensively fragmented and frequently limits confidence in the overall crosslink assignment. Electron transfer leaves DSSO largely intact and gains little analogous benefit from its cleavability. Thus, the principal analytical advantage of DSSO for MS2 mass-modification analysis is improved sequence-specific fragment evidence consistent with published results (52), which explains why HCD-based acquisition outperformed electron-transfer-based methods in this study.

### MS2.stHCD acquisition is preferred for purified DSSO-crosslinked assemblies

Using the rabbit 80S ribosome, we compared nine acquisition strategies based on different combinations of CID, HCD and ETD fragmentations. For purified assemblies of modest complexity, the combined effects of rapid acquisition and efficient DSSO backbone fragmentation favor MS2.HCD-based workflows. MS2.stHCD is a practical default because it performed similarly to single-energy MS2.HCD without a meaningful acquisition-time penalty and provides broader collision-energy coverage when diagnostic ions are also of interest. This recommendation depends partly on the analysis strategy. Search engines that require diagnostic ions must balance collision energies that favor linker cleavage with those that favor backbone fragmentation. Our collision-energy results therefore support stepped HCD when both forms of evidence are needed. HCD generated diagnostic pairs most efficiently at lower collision energies than those that maximized CSM identification, while CID was less sensitive than HCD to collision-energy variation and may also provide an effective compromise between diagnostic-ion and backbone-fragment formation. However, because we did not test a stand-alone MS2.CID strategy, the present data do not establish whether optimized solo MS2.CID would perform equivalently to MS2.stHCD. There is no compelling evidence to support acquiring separate CID and HCD scans from the same precursor.

The contribution of accompanying MS2.CID spectra also depended on sample complexity. In the ribosome dataset, approximately one-third of the URPs reported by the MS2.CID-MS3.CID workflow were contributed by the MS2.CID analysis. In the HEK293 whole-cell lysate, MS2.CID contributed only approximately 8% of the inter-protein URPs from the same combined workflow. The smaller ribosome search space permits classification with less fragment evidence, whereas proteome-wide searches require stronger sequence-specific evidence to achieve the same nominal FDR. Thus, MS2.CID spectra optimized primarily for diagnostic-ion formation add relatively little to proteome-scale identification, even though combining the MS2 and MS3 results can modestly increase total coverage. The greater sampling depth of MS2.stHCD also produced a higher rate of apparent structural-distance violations in the ribosome dataset. Many of these long-distance crosslinks were supported by high-quality spectra and were consistent with collided ribosome dimers, conformational heterogeneity, or low-abundance aggregated species rather than random incorrect assignments. This illustrates a general limitation of validating sensitive XL-MS methods against a single static structure: deeper acquisition may increase apparent violations by detecting genuine conformations or assemblies that are absent from the reference model.

### MS2.stHCD and MS3.CID provide complementary advantages at proteome scale

At proteome scale, the preferred workflow depends on the reporting objective. MS2.stHCD provides greater spectral and residue-pair depth because it samples precursors rapidly and efficiently generates informative DSSO backbone fragments. MS3.CID provides nearly comparable protein-pair coverage with much simpler linear-peptide searches and excellent control of error in crosslink reconstruction. The accompanying MS2.CID analysis added relatively little protein-pair coverage in the HEK293 dataset while reintroducing the computational and FDR-estimation demands of MS2 mass-modification searching.

MS3.CID has two major practical advantages for proteome-scale analysis. First, MS3 spectra can be searched as linear peptides, allowing large datasets to be analyzed within hours rather than the multiple days required for proteome-wide MS2 mass-modification searches. Second, MS3-level analysis produces very propagated decoy assignments and is therefore less dependent on establishing technically robust error estimation and control strategies than MS2 analysis. This reflects the stringent reconstruction of crosslinks from a limited number of independently identified linear peptides constrained by linked MS1 and MS2 information, which allows very few decoy assignments to propagate to the crosslink level. In the present HEK293 dataset, no decoy MS3 assignments propagated to the CSM, URP, or protein-pair levels, even when the initial linear-peptide thresholds were relaxed. This should be interpreted as a very low empirical decoy rate rather than proof of a true 0% FDR, but it nevertheless makes MS3 error control comparatively straightforward.

MS2 mass-modification searches have different error-propagation characteristics. The initial search permits a broad range of modification masses on an internal lysine and can retain hundreds or thousands of candidate peptide assignments. Crosslink reconstruction then considers a large number of peptide combinations whose modification masses satisfy the precursor-mass relationship. This process introduces substantially more decoy candidates than the MS3 workflow and makes accurate analysis dependent on robust post-search scoring and FDR estimation. The practical choice is therefore not simply between larger and smaller identification counts.

MS2.stHCD is attractive when the goal is maximal residue-level coverage, whereas MS3.CID alone provides nearly equivalent protein-pair coverage with faster searches and more restrictive empirical error control.

The relative proportions of intra-and inter-protein identifications also differed between the workflows. MS3.CID produced a higher percentage of inter-protein protein pairs than MS2.stHCD, but this is unlikely to reflect preferential sampling of inter-protein species. Instead, the difference may reflect differential recovery of intra-protein assignments. In MS2 mass-modification searches, a strongly assigned peptide can constrain a weakly fragmented partner to the same protein, substantially reducing the effective candidate space. By contrast, the current MS3 pipeline evaluates the two constituent peptides independently and requires both to pass the same linear peptide identification criteria before crosslink reconstruction, regardless of whether the resulting crosslink is intra-or inter-protein. Relaxing the initial MS3 peptide thresholds and applying more stringent classification after crosslink reconstruction could potentially recover additional intra-protein assignments.

### Toward acquisition strategies that combine MS2 depth with selective MS3 confirmation

Simple precursor charge, *m/z*, and peptide-length features did not identify a clear population that would selectively benefit from MS3 acquisition. More sophisticated real-time scoring, partial-sequence recognition, or firmware-level search logic will therefore likely be required. An ideal strategy would combine the acquisition depth and efficient DSSO fragmentation of MS2.stHCD with selective MS3 confirmation of spectra that cannot be resolved confidently at the MS2 level. Real-time acquisition logic could potentially trigger MS3 based on diagnostic-ion patterns, partial peptide sequences, or other features of the precursor and product-ion spectra. The REACT approach developed by the Bruce laboratory demonstrates the feasibility of selectively triggering MS3 using features specific to Protein Interaction Reporter reagents (57). However, that approach does not directly leverage a complete MS2 crosslink assignment before deciding whether additional scans are needed.

Future development of Touchstone will also focus on integrating evidence across scan types, including MS2.HCD and MS3.CID, to combine their complementary strengths within a unified classification framework.

In summary, Prospector-Touchstone provides a general and accurately calibrated platform for MS2-based analysis of cleavable and non-cleavable crosslinks. The results demonstrate that diagnostic ions are useful markers of cleavable crosslinked precursors but are less informative than sequence-specific backbone fragments for resolving peptide-pair assignments. DSSO cleavage substantially improves HCD backbone fragmentation and shifts the optimal acquisition strategy away from electron-transfer-based methods. For purified assemblies, MS2.stHCD provides the greatest overall performance, whereas proteome-scale experiments involve a practical choice between the residue-level depth of MS2.stHCD and the computational simplicity and restrictive error behavior of MS3.CID. Continued development of selective MS3 triggering and integrated MS2/MS3 scoring may ultimately combine the strongest features of both approaches.

## Experimental Procedures

### Adapting Protein Prospector to Search for Cleaved Crosslinker Fragments

The assumption in most search engines is that if there is a modification on a particular amino acid, then this modification will be retained on the fragments. A few search engines, including Protein Prospector, can also consider modifications as neutral losses, where the mass is retained on the precursor ion but absent from the fragments. However, to support DSSO crosslinker analysis, the software needs to consider two modification states on fragments (alkene and sulfene), neither of which correspond to the modification on the precursor (i.e. the mass added by the crosslinker), and not to consider fragments containing the full modification. To accommodate this, in the parameter file where one defines a modification, modifications can be specified as labile by adding a * to the end of the name; e.g. Xlink:DSSO*. The elemental compositions of the retained modification/s are defined compared to an unmodified residue, which in the case of DSSO would be C3 H2 O1 S1 and C3 H2 O1. Modifications can be indicated as ‘Full’, where only the fragments of the modification are considered, or ‘Partial’, where the intact modification as well as the modification fragments are considered.

### MS-Filter for Identifying Spectra Containing Signature Ions

MS-Filter is software that filters peak list files for the presence of specific masses, mass differences, or mass losses from the precursor. To accommodate searching for spectra with signature ions from cleavable crosslinkers the user defines two parameters: (1) the elemental composition difference between the precursor ion and the summed composition of the signature ions from the two peptides: ‘Offset Elem Comp’; (2) for cleavable crosslinkers that produce two reporter ions per peptide, the elemental composition difference between these reporter ions: ‘Delta Elem Comp’. An additional parameter added was the option to require the two reporter ions for a peptide to be seen in the same charge state (‘Match Z’). These parameters are all applied to ‘Intact Pep Pairs’.

### Touchstone description and scoring function development

Touchstone is an R library developed to facilitate automated scoring, classification, summarization, visualization and biological inference from XL-MS data. Touchstone supports analysis of cleavable and non-cleavable MS2 crosslinking data. All Touchstone functions are written in the R programming language, and Touchstone can be invoked in an interactive or scripted R environment (such as RStudio). We have also developed a web-based interface to Touchstone which runs under Shiny (https://shiny.rstudio.com). The Shiny version of Touchstone integrates with the Prospector web-based application when it is run in Pipeline mode. This requires a server configuration running both Apache and Shiny-Server. Touchstone reads Prospector-generated Search Compare results, generates a support vector machine (SVM) based scoring model to distinguish target from decoy hits, and summarizes at defined false-discovery rates and specified levels of summarization (CSMs, URPs, peptide pairs, protein pairs, or module pairs). Shiny Touchstone provides links to annotated spectra and has checkboxes that facilitate manual inspection and removal of spectra for edge-cases. Navigation to crosslinks of interest is facilitated by an interactive table (built with the DT R library) and an interactive protein-pair graph. An optional module file can be provided by the user to map defined regions of proteins to modules or domains with corresponding high-resolution structure specified in the form of a PDB code or PDB/mmCIF file. We use the term “module” to mean a stable molecular assembly (*e.g.* the Mediator head module) since many larger XL-MS studies benefit from tabulating data at this level. When PDB mappings are specified in the module file, Cα-Cα distances are measured by mapping the crosslinked residues on the appropriate PDB chain and file and using the Bio3D R package(60) to obtain the atomic coordinates and calculate Euclidean distances. For each mappable crosslink, a randomized Cα-Cα distance sampled from all possible K-K residues in the PDB set is also reported.

Convenience functions are included to annotate protein–protein interactions with external resources such as STRING-db (*e.g.*, via the getStringScores function). Shiny Touchstone generates interactive plots of protein-pairs and module-pairs which facilitate visualization and selection of crosslinks of biological interest. Touchstone can export classified or user-selected crosslinks for visualization in xiView and as ChimeraX pseudobond files. Installations configured with xiNet (61) can additionally generate interactive network views through the Shiny interface. Touchstone also includes internal R functions for plotting protein-pair and module-pair networks outside the Shiny interface.

The Touchstone SVM scoring model is trained on two mutually exclusive training sets drawn from the putative spectral matches. Each training sample defaults to 20,000 rows. For results less than 40,000 rows, the data is split in two. The model is trained on each training set independently and then used to score all data not used in training.

Scores are averaged for data points that were scored by both models. Touchstone chooses features based primarily on whether the data is detected to be a multi-protein analysis or whether there is a single target protein observed. An analysis is determined to be single protein if the number of high-scoring CSMs to the top-ranked intra-protein hit exceed the second ranked intra-protein hit by 100. For multiprotein systems, the Search Compare columns used as features are: Score Difference, Percent of TIC matched, precursor mass error, precursor charge, numURPs, numCSM, and xlinkClass. Score Difference is the difference in Prospector Score between the best crosslinked hit and the best scoring linear peptide hit and can be conceptualized as the score contributed by the worse fragmenting peptide of the crosslinked pair to the overall crosslink score. Precursor mass error is featurized as an absolute value after normalizing to the mean. NumCSM is the number of CSMs matched to a unique residue pair and serves as a rough measure of abundance. NumURPs is the number of URPs tabulated by protein pair. The logic of these last two features is that multiple unique residue pairs matching to the same protein pair are more likely to be real than random. Likewise, multiple spectral matches matching to the same URPs are more likely to be real than random. Finally, xlinkClass can be either inter-protein or intra-protein as intra-protein crosslinks are much less likely to form by random chance alone. Hence, the SVM model considers both the quality of the match between product ion spectra and theoretical peptides (Score Difference, percTICmatched, precursor mass error), as well as other features in the Search Compare output which have different distribution between correct and random matches (precursor charge state, number of CSMs per URP, number of URPs per PP, and xlinkClass). For single protein searches, the features are the same except that xlinkClass and numURPs are omitted since this leads to overfitting the data (e.g. decoys are all inter-protein and targets are all intra-protein hits). Touchstone makes use of the e1071 machine learning R library for SVM modeling.

Across multiple exploratory datasets, we observed that SVM performance was relatively insensitive to the standard cost and tolerance hyperparameters when using a linear kernel. Hence, Touchstone omits an explicit hyperparameter optimization step. Instead, performance is optimized through a pre-filtering strategy based on Score Difference thresholds. Specifically, input data are filtered at thresholds corresponding to approximate FDR levels of 45%, 35%, and 25%, in addition to the unfiltered dataset.

SVM training is performed independently at each pre-filter level, and the model yielding the highest number of confidently classified inter-protein CSMs at the target FDR is selected. This approach was found to be important for achieving robust performance, particularly in proteome-scale searches. More recent versions of Touchstone incorporate alternative kernels (e.g., radial basis functions), hyperparameter optimization, and modified feature sets; however, these were not used in the present analysis.

FDR calculation in Touchstone is based on estimating the number of target-target crosslinked pairs with one and two incorrectly identified peptides based on the number of target-decoy hits and decoy-decoy and adjusting for the increased size of the target-decoy search space hits as derived in several references(35, 62–65). We extend the same logic to decoy databases which are scaled by an arbitrary scaling constant *k* (Supplemental Figure 2). This enables Touchstone to utilize larger decoy databases to better estimate error in XL-MS experiments using purified components. The number of target-target hits with one wrong peptide is given by:

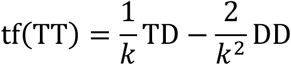

and the number with both wrong is given by:

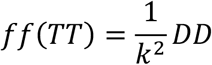

where TT, TD, and DD are the number of target-target, target-decoy, and decoy-decoy hits, and k is the scaling factor describing the ratio of the decoy database size to the target database size.

Touchstone has an option to either treat all crosslinks as belonging to a single class or to treat inter-protein and intra-protein crosslinks as separate classes for the purposes of estimating FDR and calculating acceptance thresholds. The user specifies which summarization level the FDR and acceptance thresholds should be calculated on. We agree with the conclusions of Rappsilber and co-workers that results should be classified at the summarization level of interest and that inter-protein and intra-protein crosslinks have different distributions of incorrect hits and should be treated separately in most cases(20).

### Experimental Design and Statistical Rationale

XL-MS acquisition and search strategies were evaluated using newly generated DSSO-crosslinked yeast 80S ribosome and HEK293 datasets together with comparisons to previously published benchmarking studies. Comparative analyses were primarily based on the number of accepted CSMs, URPs, and protein-pairs, identified at defined false discovery rate (FDR) thresholds. False discovery rates were estimated using target-decoy approaches implemented in Touchstone or the corresponding external software packages. Unless otherwise specified, results were filtered to 1% FDR at the indicated level of summarization. Separate intra-protein and inter-protein FDR thresholds were used for all analyses. In some classes, such as intra-protein results or MS3.CID results, not enough decoy matches were detected to make 1% FDR and the nominal error rate is below this.

Ribosome acquisition-method comparisons were performed using two technical replicate analyses acquired using the indicated fragmentation methods. Summarized crosslinking data are reported as the mean ± standard deviation across replicates.

Methods m8 and m9 had only a single replicate and no standard deviation is reported. HEK293 analyses were acquired in biological duplicate and the total number of crosslinks at the indicated summarization level were reported. Analyses are intended as matched-method comparisons in complex mixtures rather than formal statistical comparisons between biological replicates.

### Preparation and Crosslinking of 80S ribosome

Ribosomes were isolated from total Rabbit Reticulocyte Lysate via sedimentation through a 300 µl high salt sucrose cushion (0.5 M sucrose, 500 mM potassium acetate, 50 mM Hepes pH 7.4, 5 mM magnesium chloride, 0.25% digitonin) at 355,000 x g for 45 min in a TLA120.1 rotor. Pellets were resuspended in 150 mM potassium acetate, 50 mM Hepes pH 7.4, 5 mM magnesium chloride, to a concentration of 0.5 mg/ml, determined by A_260_. Crosslinking was performed by adding disuccinimidyl sulfoxide (DSSO, prepared as a fresh 10 mM stock in DMSO) to a final concentration of 0.5 mM and incubating for 30 min at 35°C. Crosslinking reactions were mixed by lightly agitating the tube every 5 min during incubation, and quenched by adding 20 mM Tris pH 8.

Reactions were TCA precipitated before processing for mass spectrometry. Pellets were washed with ice cold acetone to remove excess lipid and detergent, and then pelleted again.

Approximately 400 µg of pelleted material were reduced with TCEP, alkylated with iodoacetamide, and subjected to two rounds of trypsin digestion. Peptides were desalted on a C_18_ macrotrap column (Michrom Bioresources) and then fractionated on a size-exclusion column (Superdex Peptide, Cytiva). The four crosslinked enriched SEC fractions were analyzed as separate LC-MS acquisitions. Each SEC fraction was acquired 1 to 3 times using different MS methods (see below). Detailed protocols are available elsewhere(3).

### Preparation and Crosslinking of HEK293 Lysates

DSSO Cross-linked whole cell lysates were prepared as previously described(14). Briefly, HEK293 cells were grown to 90% confluence and ruptured by suspension in lysis buffer (100 mM sodium chloride, 50 mM sodium phosphate, 10% glycerol, 2 mM ATP, 1 mM TCEP, 10 mM MgCl2, 1x protease inhibitor (Roche), 1× phosphatase inhibitor, and 0.5% NP-40, pH 7.5). Cellular debris was removed by centrifugation and the supernatant was adjusted to 1mg/ml and crosslinked with 2 mM DSSO for 1 h at room temperature. The reaction was quenched with 50 mM ammonium bicarbonate. Cross-linked lysates were reduced, alkylated with iodoacetamide, and digested with sequential LysC and trypsin additions, using a filter-aided sample preparation protocol. The peptide digests were desalted and subjected to SEC chromatography to enrich crosslinked peptides. Two crosslinked fractions were collected and further fractionated by high-pH-reverse-phase tips (HpHt)(14). Six HpHt fractions were collected for each SEC fraction. The experiment was performed with two biological replicates, resulting in a total of 24 fractions for LC MS^n^ analysis.

### Mass Spectrometry

All LC-MS analysis was performed with an Orbitrap Fusion Lumos mass spectrometer (Thermo Scientific) coupled with a nanoelectrospray ion source (Easy-Spray, Thermo) and M-Class NanoAcquity UPLC system (Waters) with a 5 µl injection loop. Injections were made in 2.5 µl of mobile phase A, using the partial-loop mode.

Mobile phase A was 0.1% formic acid in water and mobile phase B was 0.1% formic acid in acetonitrile. DSSO crosslinked 80S ribosome fractions were separated on an Easy Spray 15 cm x 75 µm column packed with 3 µm 100Å Pep Map C_18_ resin (Thermo). The samples were loaded at 600 nl/min for 10 minutes in 5% B. The injection loop was then switched offline and the flow rate set to 300 nl/min for the duration of the acquisition. After 3 minutes of equilibration, peptides were separated with three successive linear gradients: to 10% B over 12 minutes, to 25% B over 75 minutes, and finally to 105% B over 5 minutes (total separation time: 95 min). The column was then washed at 75% B and re-equilibrated at 5% B.

Crosslinked HEK293 whole cell lysates were separated on a 50 cm x 75 µm Easy Spray column packed with 2 µm 100Å Pep Map C_18_ (Thermo). Samples were loaded at 300 nl/min for 20 minutes in 3.5% B. The loop was then switched offline and a linear gradient to 30% B at 300 nl/min over 135 minutes was executed. The column was then washed at 75% B and re-equilibrated at 3.5% B.

All MS1 scans were acquired in the Orbitrap at 120,000 resolving power. Ions were measured between 375-1500 m/z, with 30% RF Lens, Automated Gain Control (AGC) set to 4.0e5 and maximum injection time of 50 ms. The Easy-IC internal calibration was activated.

Ions were selected for MS2 analysis using monoisotopic precursor selection (MIPS) filter set to “peptide”, and intensity threshold of 5e4, an isolation window of 1.6 m/z units in the quadrupole and dynamic exclusion set to 30 sec. For 80S ribosome analysis charge states of 3-8^+^ were allowed. For HEK293 analysis, charge states of 4-8^+^ were allowed. All MS2 scans were acquired in the Orbitrap at 30,000 resolving power.

MS2-HCD: activation was performed at 28% normalized collision energy (NCE) with an AGC target of 1e5 and max injection time of 150 ms.

MS2.stHCD: was the same as MS2-HCD except using 23-28-33% NCE steps. MS2-ETD: used the charge calibrated injection time option for the reagent ions, and an AGC target of 2e5 with 200 ms max injection time.

MS2-EthcD: was identical to ETD with supplemental HCD activation at 10% NCE. MS2-CID: was similar to MS2-HCD except used 26% NCE CID activation for 10 ms.

The activation Q was 0.25 and max injection time was 100 ms. Methods based on a single acquisition method (*i.e.*, MS2-HCD) took 18 product ion scans per precursor scan. Tandem methods (*i.e.* MS2-CID-MS2-HCD) split these evenly so that each precursor was acquired by 2 of each scan type. For instance, 9-CID product ion scans and 9-HCD scans were performed for each precursor scan.

For MS3 selection, the MIPS filter was set to peptide with intensity threshold of 5e3. Charge states between 2-6 were accepted. The precursor ions were excluded from selection using a-1 to +2 m/z filter. A targeted mass filter looked for 2 precursors in each group with a delta mass of 31.9721. The partner intensity range was 10-100% relative to the most intense precursor. Partner ions were required to be in the same charge state and MS3 scans were taken on both ions in the pair. A mass tolerance of 30 ppm was used.

MS3.CID: Both the MS and MS2 isolation windows were 2 m/z units. The CID parameters were set as in MS2-CID except using 35% NCE. Selection and measurement took place in the linear ion trap using the rapid scan rate setting. The AGC target was 2e4 and maximum injection time was 100 ms.

MS2.CID-MS3.CID (method 2): an MS1 scan was taken every 5 seconds, with 4 MS3 scans taken per MS2 scan. All parameters were as above except that the MS2-CID used a 22% NCE. The method is based on Kao, *et al*(30).

MS2.CID-MS3.CID-MS2.ETD (method 3): For each MS1 scan, 9 MS2-CID scans were taken with 4 dependent MS3 scans and 9 further MS2-ETD scans on the same MS1 precursors. All parameters are as above except that the MS2-CID used a 30% NCE. The method is based on reference Liu *et al*(15).

For the HEK293 acquisition some minor changes were made. For MS2-EThcD, one biological replicate was acquired with the charge calibrated injection times activated.

However, due to a calibration error during instrument tuning, one of the biological replicates was acquired using 25 ms of ETD reaction time, a 5e4 reagent ion AGC and 25 ms max reagent ion injection time. MS2-CID-MS3-CID was the same as for the ribosome study except that the top 4 most intense MS2-CID ions were selected for MS3 rather than using the delta mass method. All the HEK293 methods used a precursor charge state range of 4-8^+^ as noted earlier.

### Data analysis

Peaklists in mgf format were generated using PAVA, our in-house tool which serves as an interface to Thermo’s RawFileReader DLL. PAVA can either use the precursor charge-state and m/z reported by RawFileReader or it can utilize Monocle(66) to recalculate the monoisotopic peak. We used the Monocle calculated values for MS2 peaklists, but not MS3 peaklists. PAVA generates separate mgf files for each type of acquisition. For instance, MS3-CID, MS2-CID, MS2-EThcD, spectra are all written as separate peaklists from data acquired using Method 3. Each set of peaklists were searched separately and results combined downstream as detailed below. All MS2-data were searched using Protein Prospector (version numbers between 6.3.23 to 6.4.5).

Peaklists were searched using the appropriate instrument type: ESI-Q-high-res for MS2-HCD and MS2-CID, ESI-ETD-high-res for MS2-ETD, ESI-EThcD-high-res for MS2-EThcD, and ESI-ION-TRAP-low-res for MS3-CID data. All searches used Trypsin as the digestion enzyme, carbamidomethylation of Cys as a fixed modification, up to 3 variable modifications per peptide including: incorrect monoisotopic mass assignment (neutral loss of 1Da) and methionine oxidation. A maximum of 100 peaks were considered for database matching. Other parameters varied between analyses:

### Synthetic Cas9 benchmark library

The target sequence database was as used in the publication and consisted of the Cas9 sequence from *S. pyogenes* plus an additional 116 sequences of common contaminants drawn from the Crapome database. Various randomized databases were appended for the decoy database as detailed in the text. The search parameters followed Beveridge *et al*(44) as closely as possible. 3 missed cleavages were allowed. The crosslinking reagent was set to DSS with a mass modification range of 500 to 6000 Da and 1000 saved intermediates. Additional variable modifications were Gln to pyroglutamate formation at a peptide N-terminus and dead-end modification by DSS at the protein N-terminus or on uncleaved lysine residues. The precursor and product ion mass tolerances were set to 5 ppm and 20 ppm respectively.

### Synthetic ribosome benchmark library

We utilized the target sequence databases distributed with the manuscript consisting of 171 sequences from *E. coli* (including the ribosomal sequences) concatenated with either 0, 500, 5000, or 20163 additional human sequences. The searches performed against the base version of the main DSSO library (*e.g.* with no additional human proteins) used a decoy database consisting of 171 random sequences which were each 10x as long as the target sequences (“10x”). The searches performed against the main library with HEK293 digests spiked in utilized a decoy database of made from scrambled version of the target sequences with the same number and size of sequences (“1x”). The search parameter were matched as closely as possible to Matzinger *et al*(45). 3 missed cleavages were allowed and the precursor and product ion tolerance were set to 5 and 10ppm. The crosslinking reagent was DSSO* with mass modification range between 600-4000 Da and 1000 saved intermediates. Dead-end modification by cleavable DSSO* at the N-terminus of the protein and uncleaved lysine residues were considered as variable modifications in addition to the base ones mentioned above.

### Rabbit ribosome searches

The target database of 77 rabbit ribosomal subunits was taken from the structure file deposited at the protein databank (pdbid: 6hcj). Note that these sequences are truncated and residue numbering will differ from reference sequences in UniProt. The decoy database was a 10x randomized version of the target database. Two missed cleavages were allowed and precursor/product ion tolerances were set to 15 and 25 ppm. The crosslinking reagent was DSSO* (cleavable DSSO) for HCD and CID-type searches and DSSO (non-cleavable DSSO) for ETD and EThcD searches. The mass modification range was 450-4500 Da with 1000 saved intermediates. Additional variable modifications included: acetylation at the protein N-terminus, loss of the protein N-terminal methionine, glutamine to pyro-glutamate conversion at the peptide N-terminus, and dead-end modification at the protein N-terminus and uncleaved lysine residues by either DSSO or DSSO* depending on the activation type.

### HEK293 whole cell lysate searches

The human proteome was searched consisting of 20,375 sequences from the SwissProt database (accession date: Nov 2, 2020) plus a randomized 1x decoy version of this database. One missed cleavage per peptide was allowed and the precursor and product ion tolerances were 15 and 25 ppm. The crosslinking reagent was DSSO* (cleavable DSSO) for HCD and CID searches and DSSO (non-cleavable DSSO) for ETD and EThcD searches. The mass modification range was 600-4000 Da with 500 saved intermediates. Variable modifications were the same as the rabbit ribosome experiments.

### MS^3^-based Cross-link Identification

Monoisotopic masses of parent ions and corresponding fragment ions, parent ion charge states, and ion intensities from MS^3^ spectra were extracted using PAVA(14). MS^3^ data were subjected to database searching using Batch-Tag within Protein Prospector (v. 6.3.5). DSSO crosslinked ribosome searches were conducted against the same database used in the MS2 analyses, whereas a database consisting of a normal human proteome (SwissProt.2022.01.19; 20,360 entries) and randomly generated 1x decoy proteome were used for DSSO crosslinked HEK293 cell lysates. For all MS^3^ searches, the mass tolerances for parent ions and fragment ions were set as ± 20 ppm and 0.6 Da, respectively. Trypsin was set as the enzyme with three maximum missed cleavages allowed. Cysteine carbamidomethylation was set as the fixed modification. A maximum of three variable modifications were also allowed, including protein N-terminal acetylation, methionine oxidation, and N-terminal conversion of glutamine to pyroglutamic acid. In addition, two defined modifications on uncleaved lysines and free protein N termini were also selected: alkene (A: C_3_H_2_O, +54 Da) and unsaturated thiol (T: C_3_H_2_OS, +86 Da), corresponding to DSSO remnant moieties. MS^3^ search results were exported and integrated with MS^1^ and MS^2^ data using the in-house script xl-Tools to identify crosslinked peptides(14, 30).

Prospector search results were classified at the desired FDR and summarization level using Touchstone. Results were reported using separate classification thresholds for inter-protein and intra-protein links, except with the Cas9 benchmarking study (in which only one target “protein” was present). For acquisition methods that combined multiple stages of dissociation, all of the peaklists from a given dissociation and MS-level were analyzed separately. When the data were summarized to URPs or PPs, the CSM with the highest SVM score was retained across the different dissociation methods. When one of the methods was MS3-CID (for which no SVM scoring model is constructed), the MS3-CID hit was always retained over any MS2 level results. For the 80S ribosome study, all numbers are reported as the average of two technical replicates which were analyzed separately, with the exception of methods 8 and 9 for which only one technical replicate was available. For the HEK293 WCL study, two biological replicates were performed and analyzed separately but the union of the two replicates is reported. Currently, the XLtools pipeline reports each putative residue pair or homologous protein whereas the Prospector-Touchstone pipeline picks a single residue/protein pair to report. To harmonize the results for proper comparison, XLtools output was grouped by spectral identifier and a single unique result was reported if there were redundant hits. The hit was chosen to be the same as the Prospector-Touchstone output in the MS2 level analysis when possible.

## Software and Data availability

Annotated spectra matching unique crosslinked residue pairs are available on MS-Viewer (https://msviewer.ucsf.edu/cgi-bin/msform.cgi?form=msviewer) using the following Search Keys:

Rabbit 80S Ribosome DSSO crosslinks:

Method 1: lzydisfdmz

Method 2 (MS2 results): cf7pwdhhsd

Method 2 (MS3 results): m47pxy7e9z

Method 3 (MS2 results): f4usjgzlbo

Method 3 (MS3 results): jag5mqvldf

Method 4: e9xynxh8vo

Method 5: ynwzlcufu7

Method 6: t91xgoxhhe

Method 7: ytjupoge7z

Method 8: camexcdgmz

Method 9: szuwjfnwt0

### HEK293 DSSO crosslinks

MS2.HCD: zi5wgplaqa

MS2.EThcD: 8vfkyyekxk

MS2.CID: s5jdx3pthu

MS3.CID: 1bpcjgqaor

Raw MS files and peaklists are deposited on ProteomeXchange (PXD078752) via the Massive Repository (MSV000101938;).

The Touchstone R package is available on github: https://github.com/mtrnka/Touchstone

A demonstration version of the Shiny app is available at: https://prospts.shinyapps.io/tstoneapp/

## Supplemental Data statement

This article contains supplemental data.

## Acknowledgments

For work done at the UCSF Biomedical Mass Spectrometry and Proteomics Resource Center (Prof. A. L. Burlingame, Director), we gratefully acknowledge the support of the Adelson Medical Research Foundation and the University of California, San Francisco Program for Breakthrough Biomedical Research. Work performed at UC Irvine (R35GM145249) and the University of Chicago (R01GM130051) was supported by the NIH.

## Author Contributions

**MJT:** Conceptualization, Formal analysis, Investigation, Methodology, Software, Validation, Visualization, Writing – original draft, Writing – review and editing.

**CY:** Investigation, Formal analysis, Methodology, Writing – review and editing.

**PB:** Methodology, Software.

**PTM:** Investigation, Resources.

**FJ:** Investigation, Resources.

**RJK:** Supervision, Resources.

**ALB:** Supervision, Funding acquisition.

**RC:** Conceptualization, Software, Writing – review and editing.

**LH:** Conceptualization, Supervision, Funding acquisition, Resources, Project administration, Writing – review and editing.

## Declaration of generative AI and AI-assisted technologies

During the preparation of this work, the authors used ChatGPT (OpenAI) to assist with editing for clarity, concision, organization, and scientific writing. Additionally, an image in the graphical abstract was AI-generated. AI was not used for data analysis or scientific interpretation. The authors reviewed and edited all AI-assisted content and take full responsibility for the content of the manuscript.

## Abbreviations

AGC: automatic gain control
BS3: bis(sulfosuccinimidyl)suberate
CID: collision-induced dissociation
CSM: crosslinked spectral match
DSS: disuccinimidyl suberate
DSSO: disuccinimidyl sulfoxide
ETD: electron transfer dissociation
EThcD: electron-transfer/higher-energy collision dissociation
FDR: false discovery rate
HCD: higher-energy collisional dissociation
HEK293: human embryonic kidney 293 cells
LC-MS: liquid chromatography mass spectrometry
MGF: Mascot generic format
MS: mass spectrometry
MS2: tandem mass spectrometry
MS3: multistage mass spectrometry
PPI: protein–protein interaction
PTM: post-translational modification
SEC: size-exclusion chromatography
stHCD: stepped higher-energy collisional dissociation
SVM: support vector machine
TCEP: Tris(2-carboxyethyl)phosphine
TIC: total ion current
URP: unique residue pair
UPLC: ultra performance liquid chromatography
XL-MS: crosslinking mass spectrometry

