## Supplemental Tables and Figures for "Optimization of Search and Acquisition Strategies for DSSO Crosslinking Mass Spectrometry using Protein Prospector and Touchstone"

**Supplemental Table 1 – MS acquisition strategies used in this study**

| Method | Name | Ref |
|---|---|---|
| m1 | ms2.hcd-ms2.ethcd | McGilvray et al, eLife 2020 |
| m2 | ms2.cid-ms3.cid | Kao et al, MCP 2011 |
| m3 | ms2.cid-ms3.cid-ms2.etd | Liu et al, Nature Comm 2017 |
| m4 | ms2.etd | |
| m5 | ms2.ethcd | |
| m6 | ms2.hcd | |
| m7 | ms2.sthcd | Stieger et al, J Proteome Res 2019 |
| m8 | ms2.cid-ms2.ethcd | Liu et al, Nature Methods 2015 |
| m9 | ms2.cid-ms2.hcd | Ser et al, J Proteome Res 2019 |

### Supplemental Table 2

**Interprotein Ribosome putative crosslinked spectra.**
**Method 8: MS2.CID – MS2.EThcD acquisition**

| | | % Spectra containing diagnostic ion pair | | |
|---|---|---|---|---|
| **Classified*** | **Decoy** | **CID** | **EThcD** | **EThcD cross†** |
| accepted | Target | 71.7% | 11.7% | 37.1% |
| accepted | Decoy | 81.5% | 22.1% | 9.2% |
| accepted | DoubleDecoy | 11.1% | | 8.3% |
| rejected | Target | 19.8% | 6.7% | 7.7% |
| rejected | Decoy | 15.3% | 6.4% | 2.6% |
| rejected | DoubleDecoy | 1.7% | 0.1% | 0.7% |

* Met acceptance criteria for 1% FDR CSM threshold
† *EThcD MS2 spectra with diagnostic pairs detected in the linked CID MS2 spectrum*

#### Supplemental Table 3

**Interprotein Ribosome putative crosslinked spectra.**
**Method 9: MS2.CID – MS2.HCD acquisition**

| | | % Spectra containing diagnostic ion pair | | |
|---|---|---|---|---|
| **Classified*** | **Decoy** | **CID** | **HCD** | **HCD cross†** |
| accepted | Target | 73.6% | 17.4% | 38.0% |
| accepted | Decoy | 77.9% | 28.3% | 9.8% |
| accepted | DoubleDecoy | 14.3% | 33.3% | |
| rejected | Target | 21.7% | 6.5% | 4.8% |
| rejected | Decoy | 16.2% | 5.1% | 1.9% |
| rejected | DoubleDecoy | 1.3% | 2.2% | 0.2% |

* Met acceptance criteria for 1% FDR CSM threshold
† *HCD MS2 spectra with diagnostic pairs detected in the linked CID MS2 spectrum*

**Supplemental Figure 1.**

A) Prospector-Touchstone XLMS data processing pipeline. B) Screen captures from the interactive
B) Shiny version of Prospector-Touchstone - Classification Tab, C) Crosslink Table Tab,
C) D) MS-Product spectral annotation.

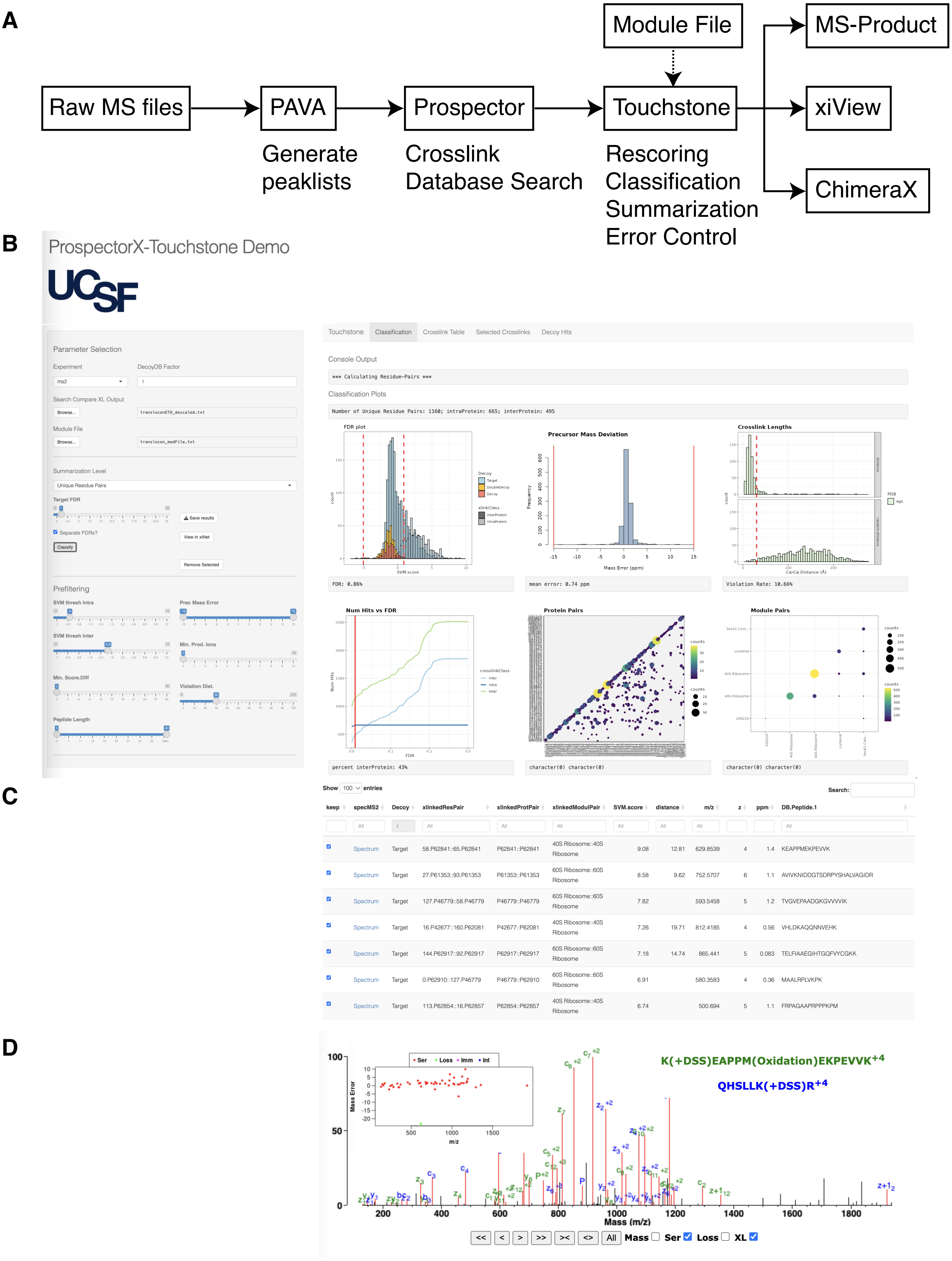

#### Supplementary Figure 2

Decoy database construction strategies. **A) Left-panel:** peptide length distributions under different methods. **Right-panel:** Example target protein and four different decoy versions. **B)** CLMS FDR estimation for arbitrarily scaled (eg 10x) decoy databases.

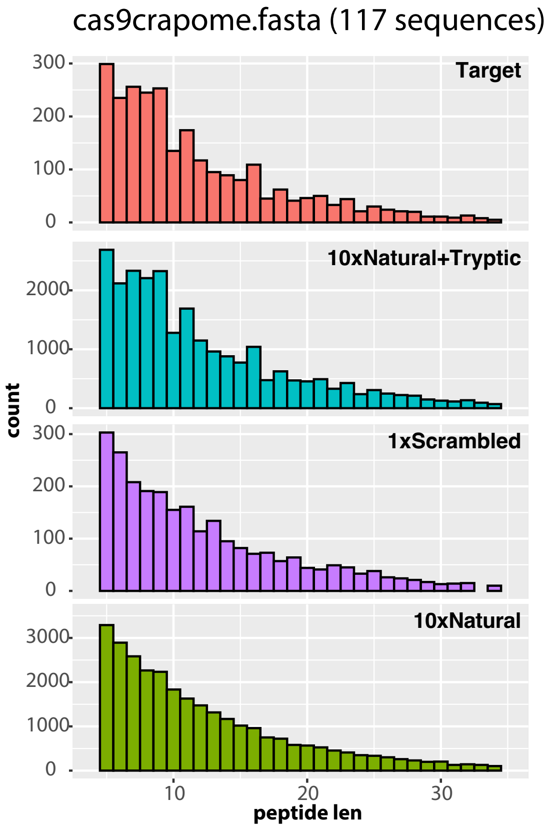


##### Target
>Protein.1
MPGHLSIKLPGKVNTREEF

##### 1xScrambled
>Protein.1
VKTHNRPPEFGGLKIMSLE

##### 1xNatural
>Protein.1
MAVSPLSSGWGFRVLLTGL

##### 10xNatural+Tryptic
>Protein.1
MLSLNYLKCPYKSAWRFLF
AYVSETSKLTLRLCSKPSW
LIGVPLGKLGPKEVGKIVA
VVLGVVFKESDRVVLRIVI
GPDSSQFKEQDKFVERVCD
NTIQDMNRESVRLSSKLEL
LEGFSVEKLWNKFLSRLYP
AINLDPHKCAVRPALKDFS
AVGELEPRVLPKDVFKDPV
QSENGYLKDSARYEWKSAP

##### 10xNatural
>Protein.1
MRQTIGPVRGRWFEANEAL
LEAFSIGYFEYLARLRCLA
AGRAEDESSAESTGCLRMT
QYIEKITTTRDRMTGQCKI
ENPIGKWSEQFQGWVPPVA
RDGSQGEVLRALFASALEL
TTSYGFRDQPVRGKDGTGL
EWPCQAWDCGSQFCNGDAR
MTGAEVPILLGSGSQQVHS
RDYDTTRYLAPGTCCVLVS

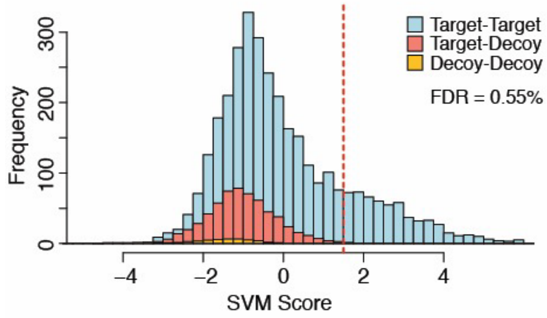


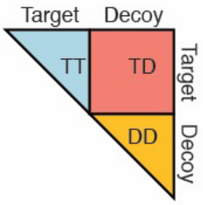


$$tf(TT) = TD - 2DD$$

$$ff(TT) = DD$$

Walzthoeni *et al, Nature Methods* 2012
Yang *et al, Nature Methods* 2012
Trnka *et al, Mol Cell Proteomics* 2014
Wang *et al, Mol Cell Proteomics* 2014
Fischer & Rappsilber, *Anal Chem* 2017

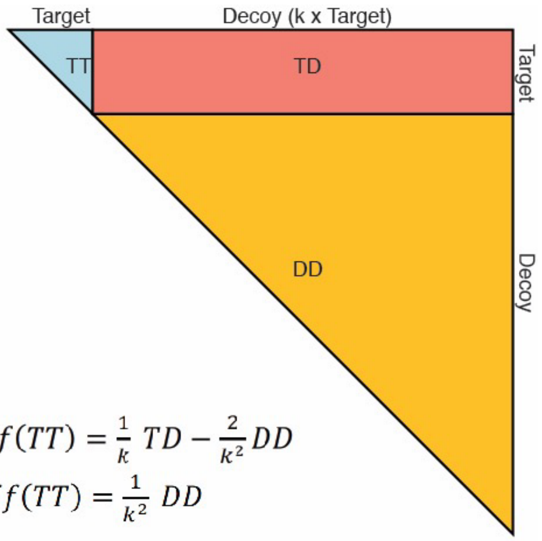


$$tf(TT) = \frac{1}{k}\,TD - \frac{2}{k^2}DD$$

$$ff(TT) = \frac{1}{k^2}\,DD$$

**Data classified at 0.5% FDR crosslinked residue pairs (merged HCD/ETD data)**

**Supplementary Figure 3**
Benchmarking Prospector / Touchstone against other MS2 based search engines. Cas9 + crapome search. ROC curves for CSM and URP-level classification. AUC values are noted in the legend.

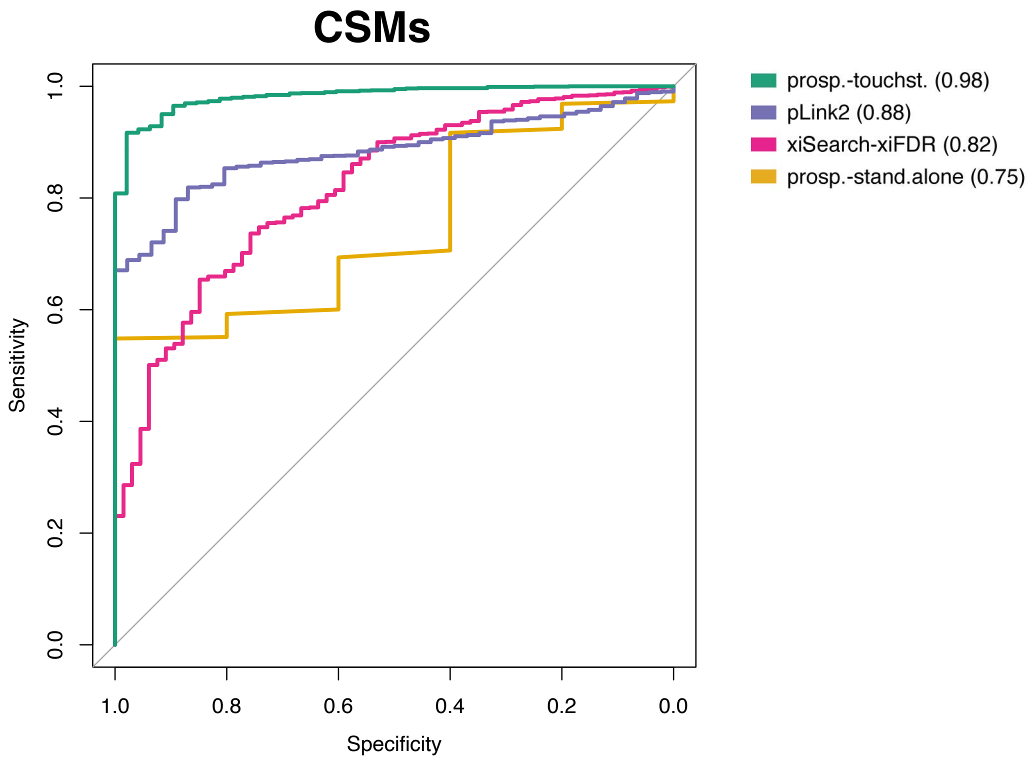


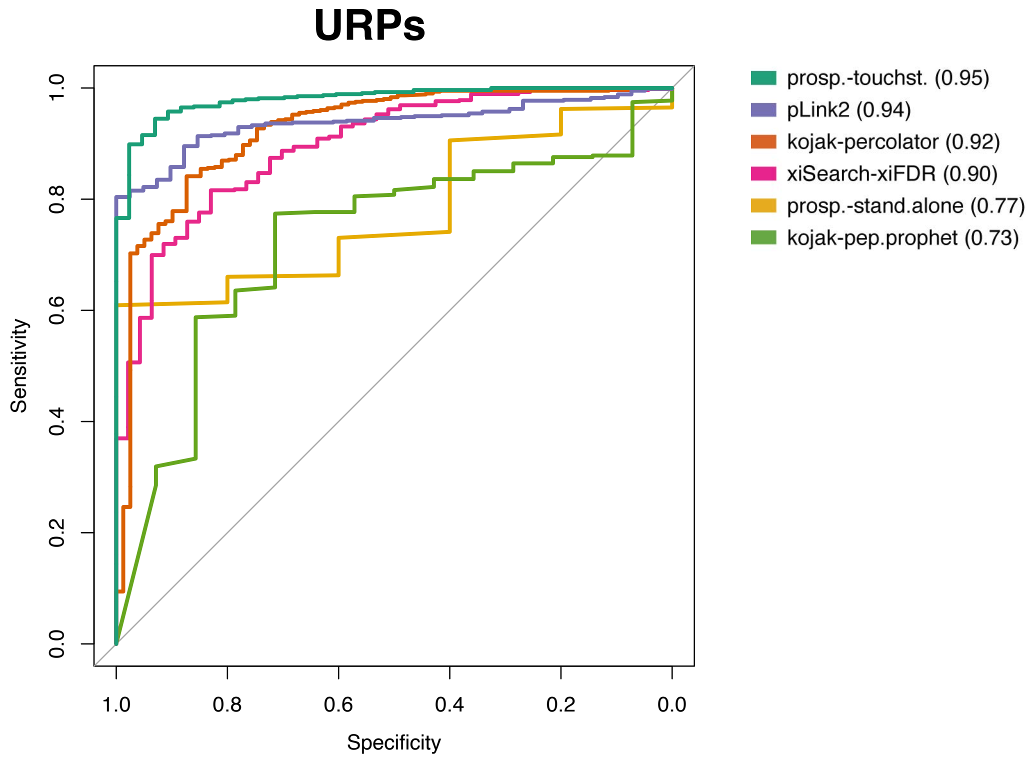

DSSO, rabbit Ribosome, stepped-HCD

DSSO* vs DSSO

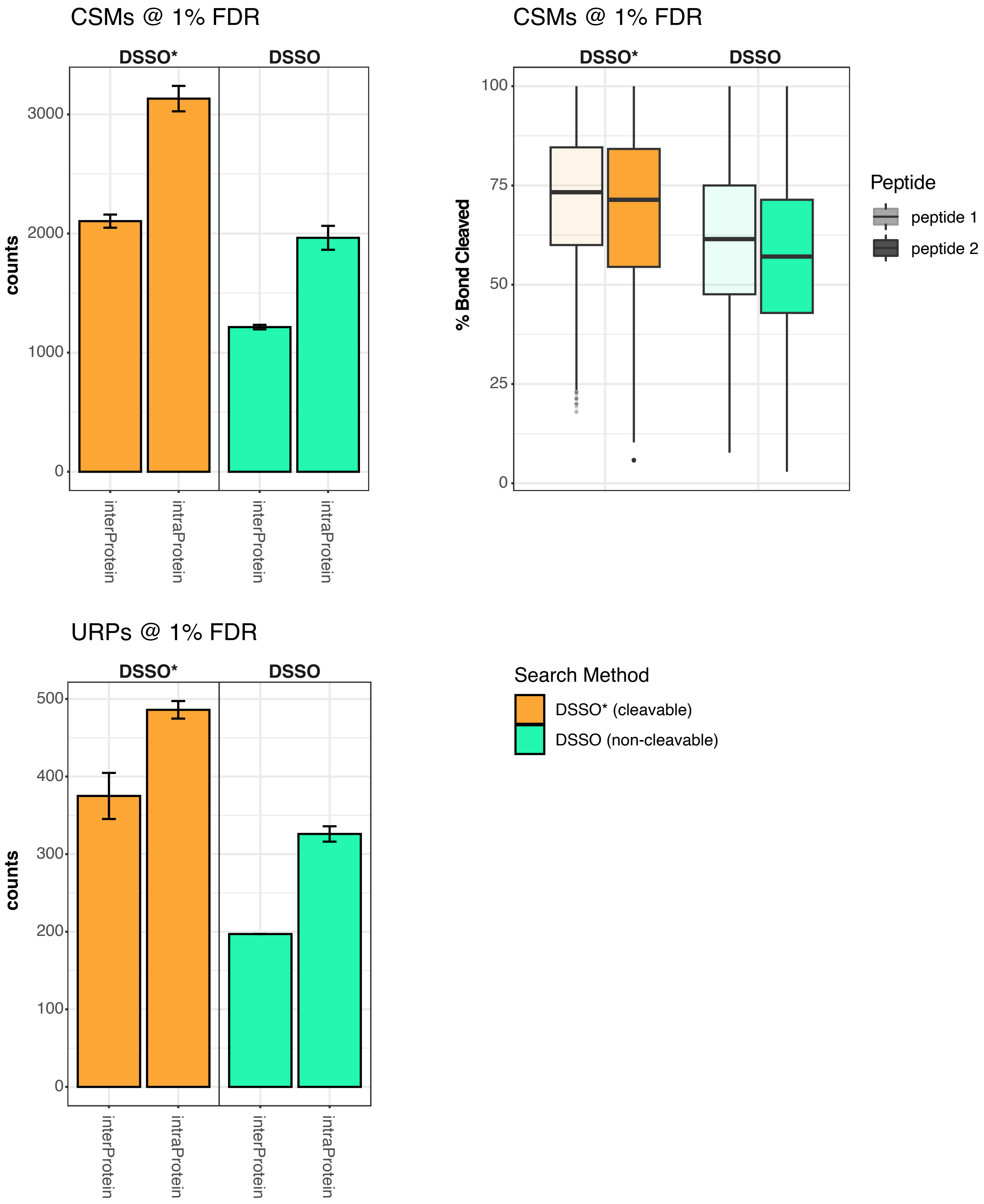


**Supplemental Figure 4.**
Stepped-HCD data from 80S Ribosome XLMS experiment searched with crosslinking reagent parameter set to either DSSO*, which treats product ions as dissociating completely at the sulfoxide, versus DSSO, which treats product ions as intact at the sulfoxide bridge as observed in electron-transfer dissociation. The DSSO option models BS3 crosslinking in which product ions spanning the site of modification are rarely observed.

**Supplementary Figure 5.**

A) Negative stain electron microscopy of 80S rabbit ribosome after 500 $\mu$M DSSO crosslinking. Examples of aggregated particles are indicated with a red arrow.

B) Western blotting of small ribosomal subunit S16 and large subunit L9 shows disappearance of monomer band and corresponding shift to crosslinked 80S band after 500 $\mu$M DSSO treatment.

**A)**

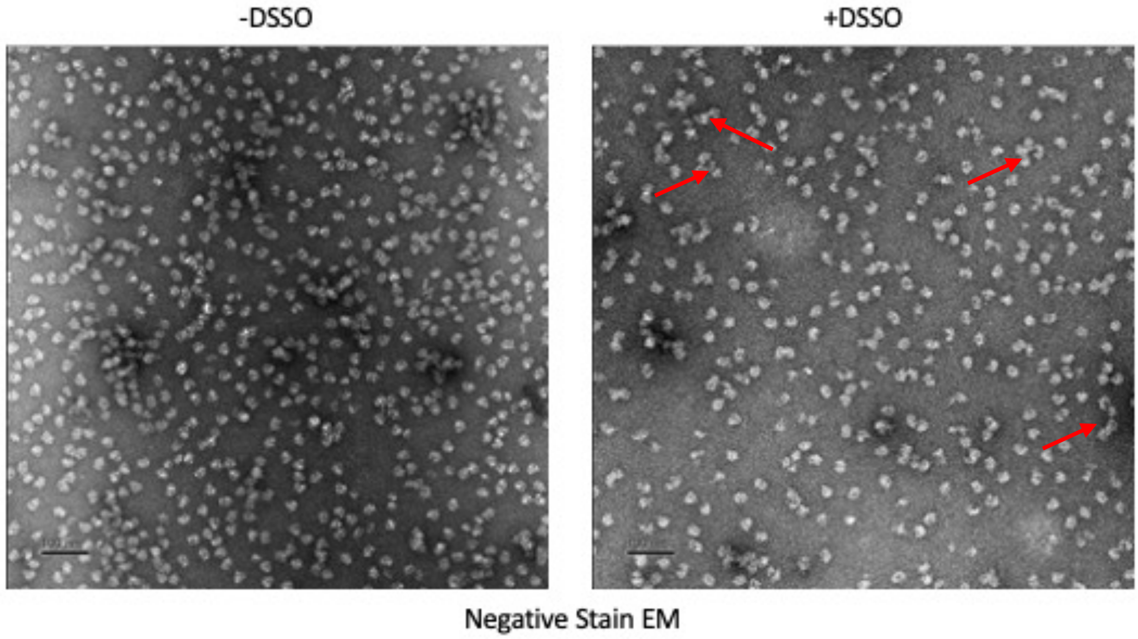


**B)**

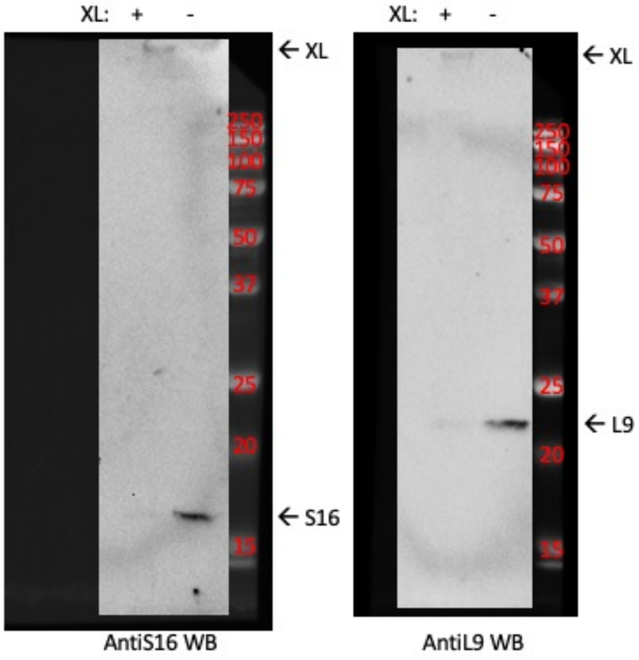

**Supplementary Figure 6.**
EThcD analysis of non-cleavable BS3 crosslinks provides higher and more symmetrical fragmentation coverage of both peptides than HCD analysis. HCD shows median product ions identifying 75% of bond cleavages from peptide 1 (more strongly fragmented peptide) and 57% from peptide 2, while median bond fragmentation observed from EThcD are 80% for peptide 1 and 75% for peptide 2. Data from McGilvray *et al* 2020.

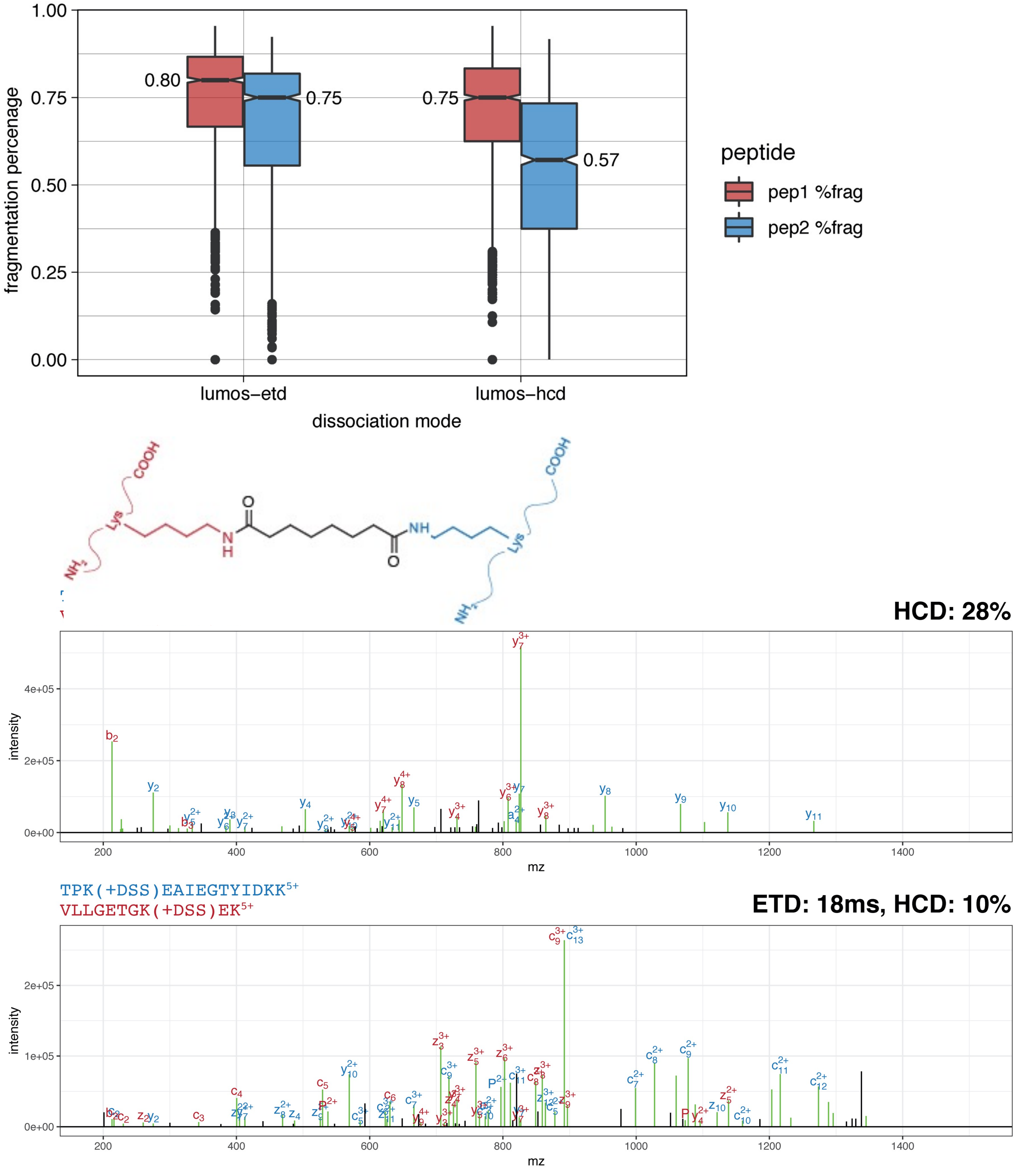

**Supplemental Figure 7.**
High quality HCD product ion spectral matches to over-length ribosomal crosslinks.

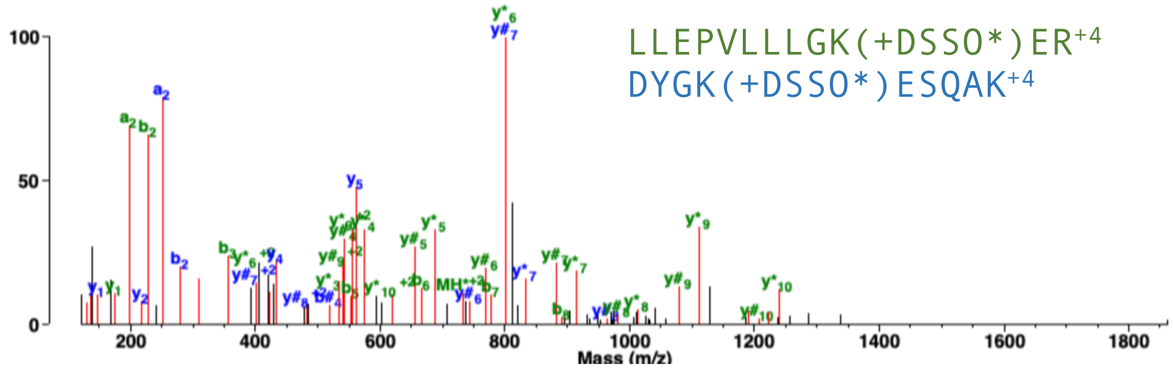


| Acc # | Prot. | Res | Score | %-Backbone ions |
|---|---|---|---|---|
| R2(G1SGX4) | RPS16 | K60 | 70.7 | 82% |
| N2(G1SFR8) | RPS12 | K116 | 86.7 | 88% |

Cα-Cα Distance: 118Å (monomer) - 24Å (dimer). 75% TIC matched.

---

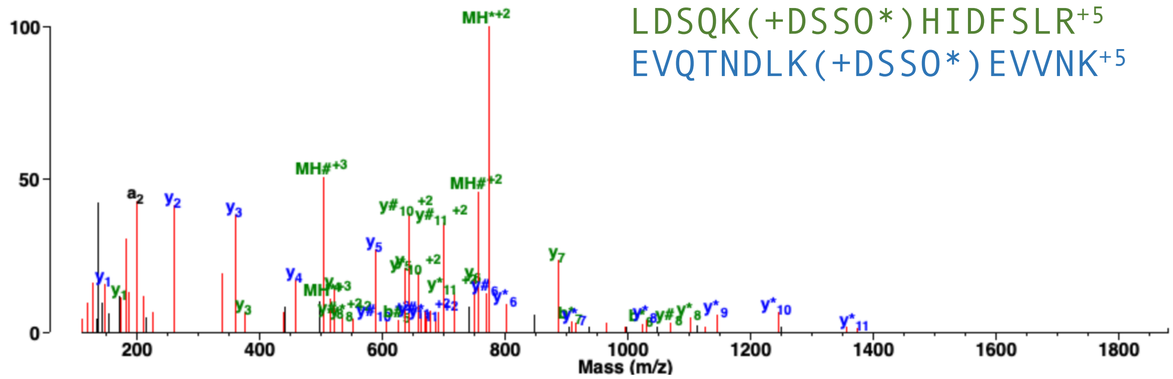


| Acc # | Prot. | Res | Score | %-Backbone ions |
|---|---|---|---|---|
| K2(B7NZS8) | RPS9 | K155 | 54.6 | 82% |
| C2(G1SS70) | RPS3A | K182 | 51.0 | 92% |

Cα-Cα Distance: 156Å (monomer) - 20Å (dimer). 83% TIC matched.

---

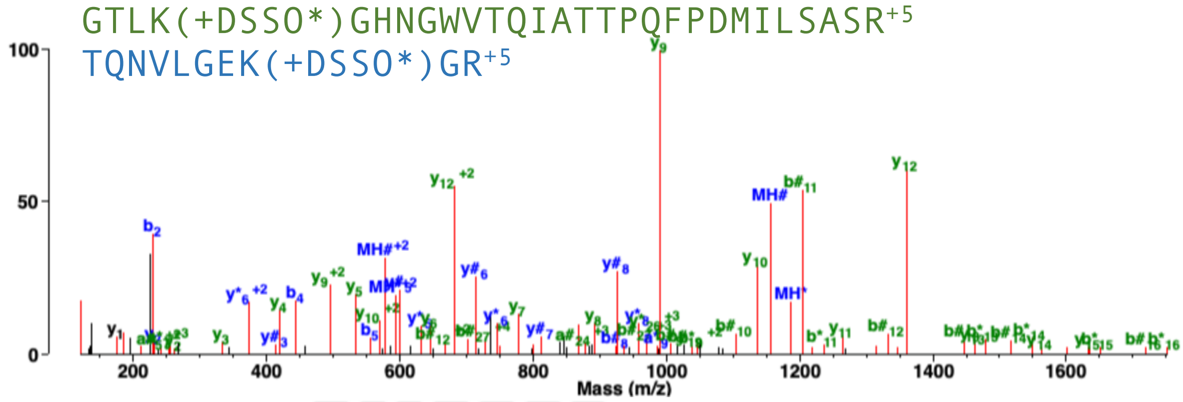


| Acc # | Prot. | Res | Score | %-Backbone ions |
|---|---|---|---|---|
| h2(G1SJB4) | RACK1 | K12 | 86.7 | 70% |
| E2(G1TNM3) | RPS3 | K62 | 33.4 | 79% |

Cα-Cα Distance: 90Å (monomer)- 21Å (dimer). TIC matched:87%

**Supplemental Figure 8.**
High quality HCD product ion spectral matches to over-length ribosomal crosslinks mapped to di-ribosome structure. Blue: nucleic acids. Purple/Green: Ribosome monomers. Darker shades indicate 40S Ribsoome. Putative crosslink highlighted in red/orange.

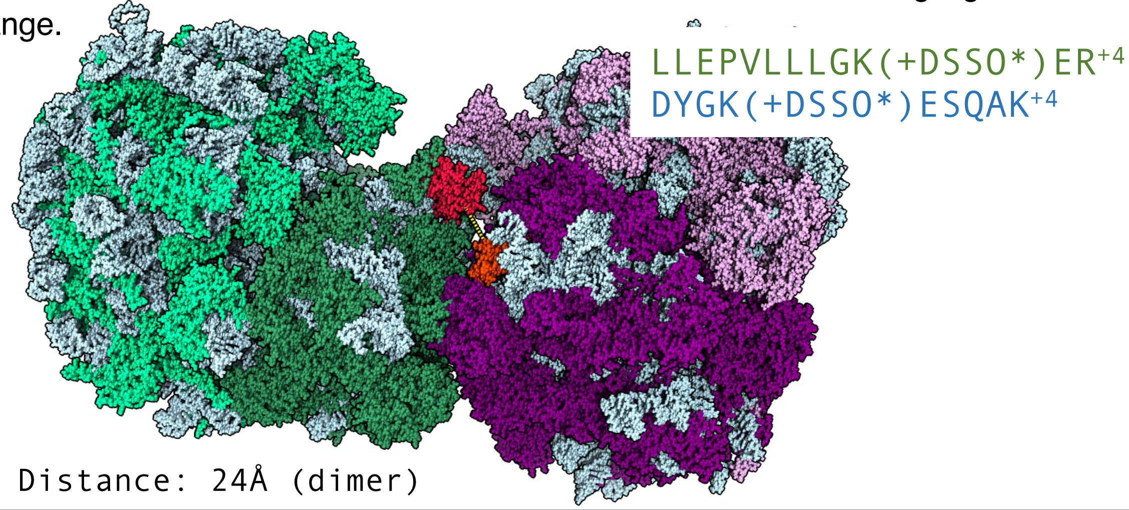


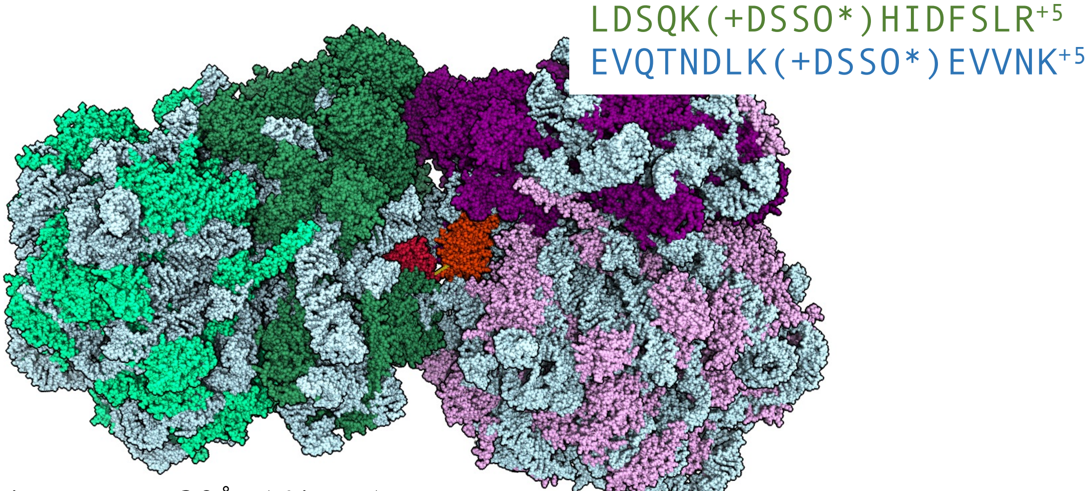


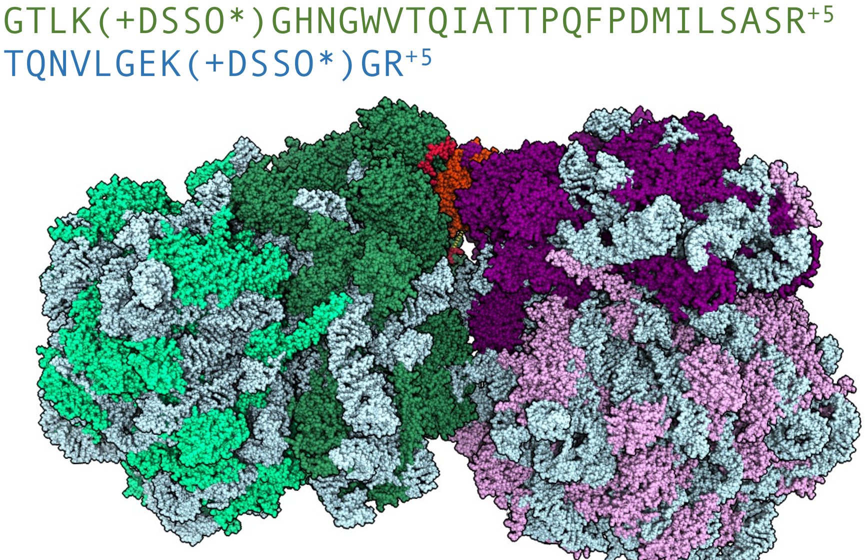

**Supplementary Figure 9.**
MS-Filter contains new options to detect spectra in a peaklist containing diagnostic pair relationships from cleavable crosslinking reagents.

For DSSO, spectra containing ions that sum to the precursor plus an offset composition ($H_1O_1$) and plus or minus a delta elemental composition ($S_1$) are detected and can be retained (equations 1-3). Product ions can be optionally be required to have the same charge state and +1 and +2Da errors can be tolerated to account for misannotated monoisotopic peaks.

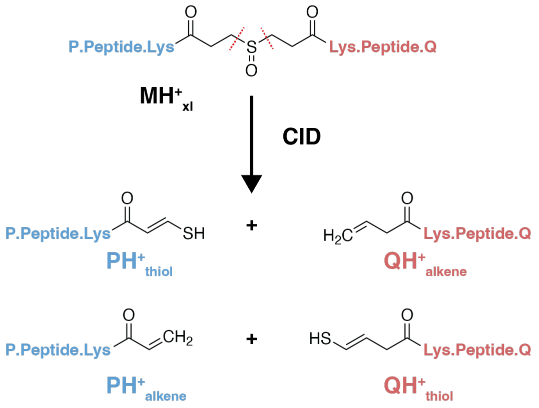


**Precursor - Product Relationships (Neutral):**

$M_{xl} = P_{thiol} + Q_{thiol} + H_2O - S_1$

$M_{xl} = P_{alkene} + Q_{thiol} + H_2O$

$M_{xl} = P_{alkene} + Q_{alkene} + H_2O + S_1$

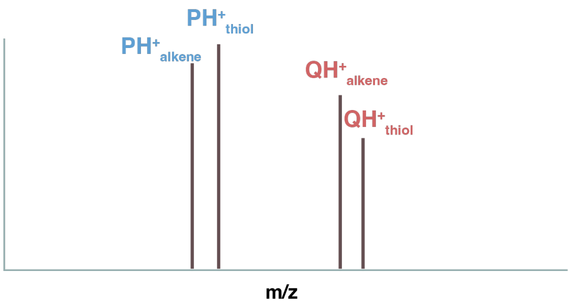


**Precursor - Product Relationships (Protonated Ions):**

1. $MH^+_{xl} = PH^+_{thiol} + QH^+_{thiol} + OH - S_1$
2. $MH^+_{xl} = PH^+_{alkene} + QH^+_{thiol} + OH$
3. $MH^+_{xl} = PH^+_{alkene} + QH^+_{alkene} + OH + S_1$

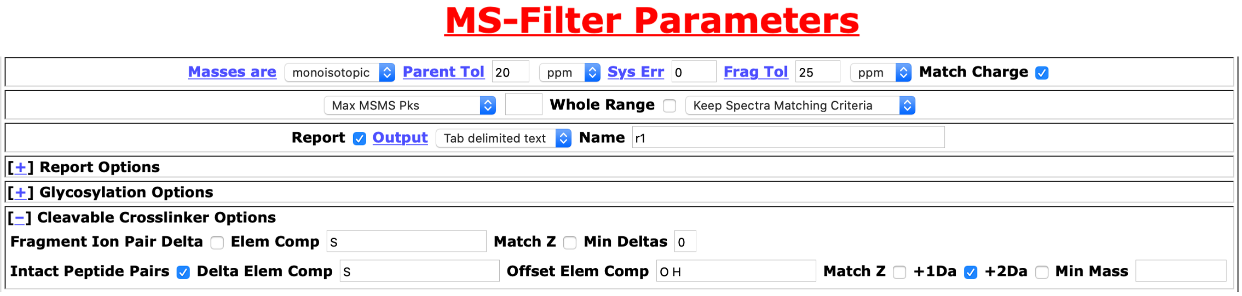
MS-Filter Parameters

Masses are monoisotopic Parent Tol 20 ppm Sys Err 0 Frag Tol 25 ppm Match Charge

Max MSMS Pks Whole Range Keep Spectra Matching Criteria

Report Output Tab delimited text Name r1

[+] Report Options

[+] Glycosylation Options

[-] Cleavable Crosslinker Options

Fragment Ion Pair Delta Elem Comp S Match Z Min Deltas 0

Intact Peptide Pairs Delta Elem Comp S Offset Elem Comp O H Match Z +1Da +2Da Min Mass

**Supplemental Figure 10.**
Optimization of Lumos collision energy (NCE) for formation of DSSO diagnostic ion pairs and Crosslinked Spectral Matches (CSMs @ 1% FDR).

DSSO crosslinked rabbit 80S Ribosome was acquired at multiple CID and HCD collision energies and A) analyzed by MS-Filter to count numbers of spectra with Thiol-Thiol, Thio-Alkene, or Alkene-Alkene pairs (left panel). B) CSMs classified at 1% FDR are reported in the right panel.

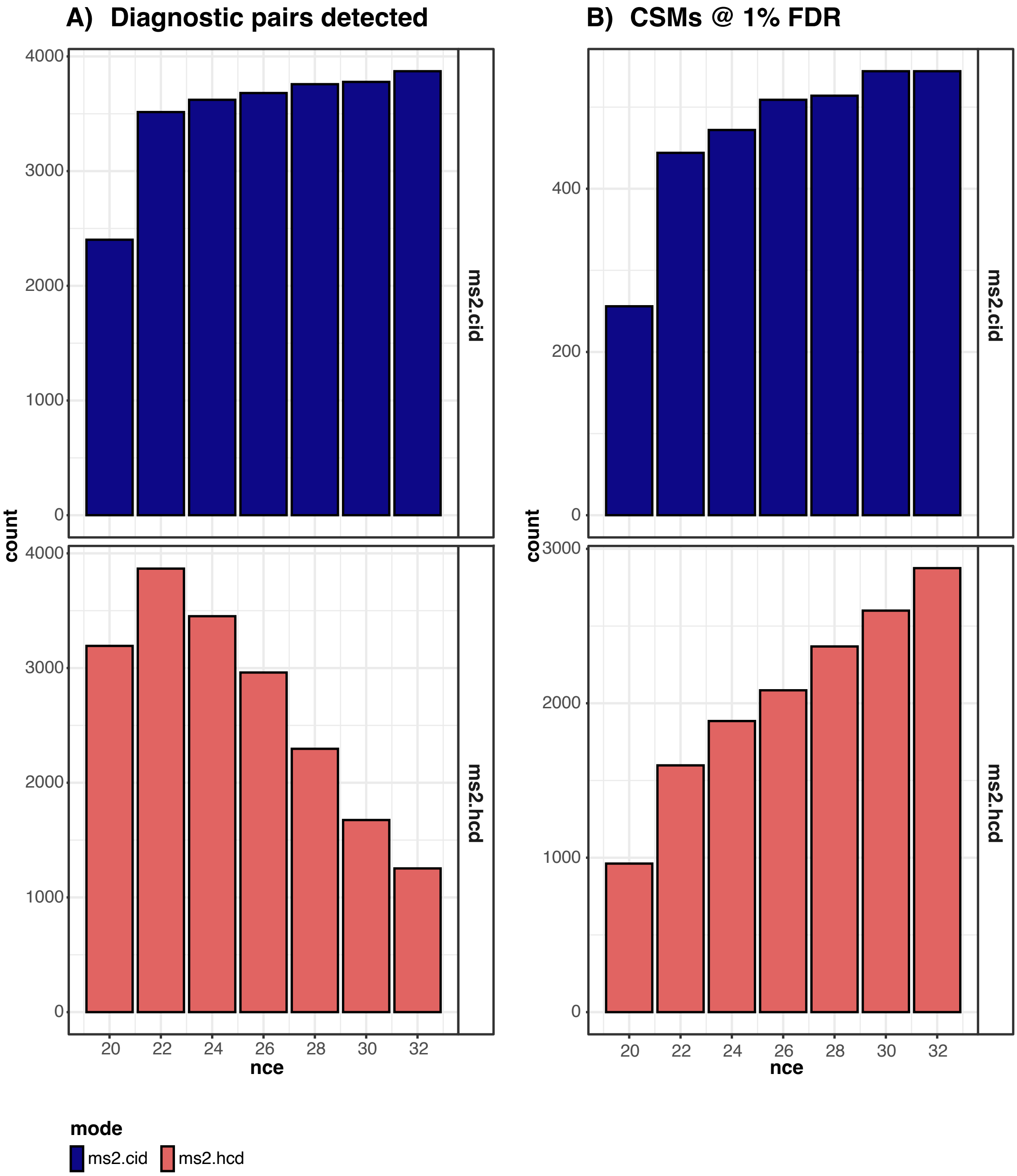

**Supplemental Figure 11**
MS2 putative inter-protein CSMs M9-CID-HCD
DSSO, rabbit Ribosome

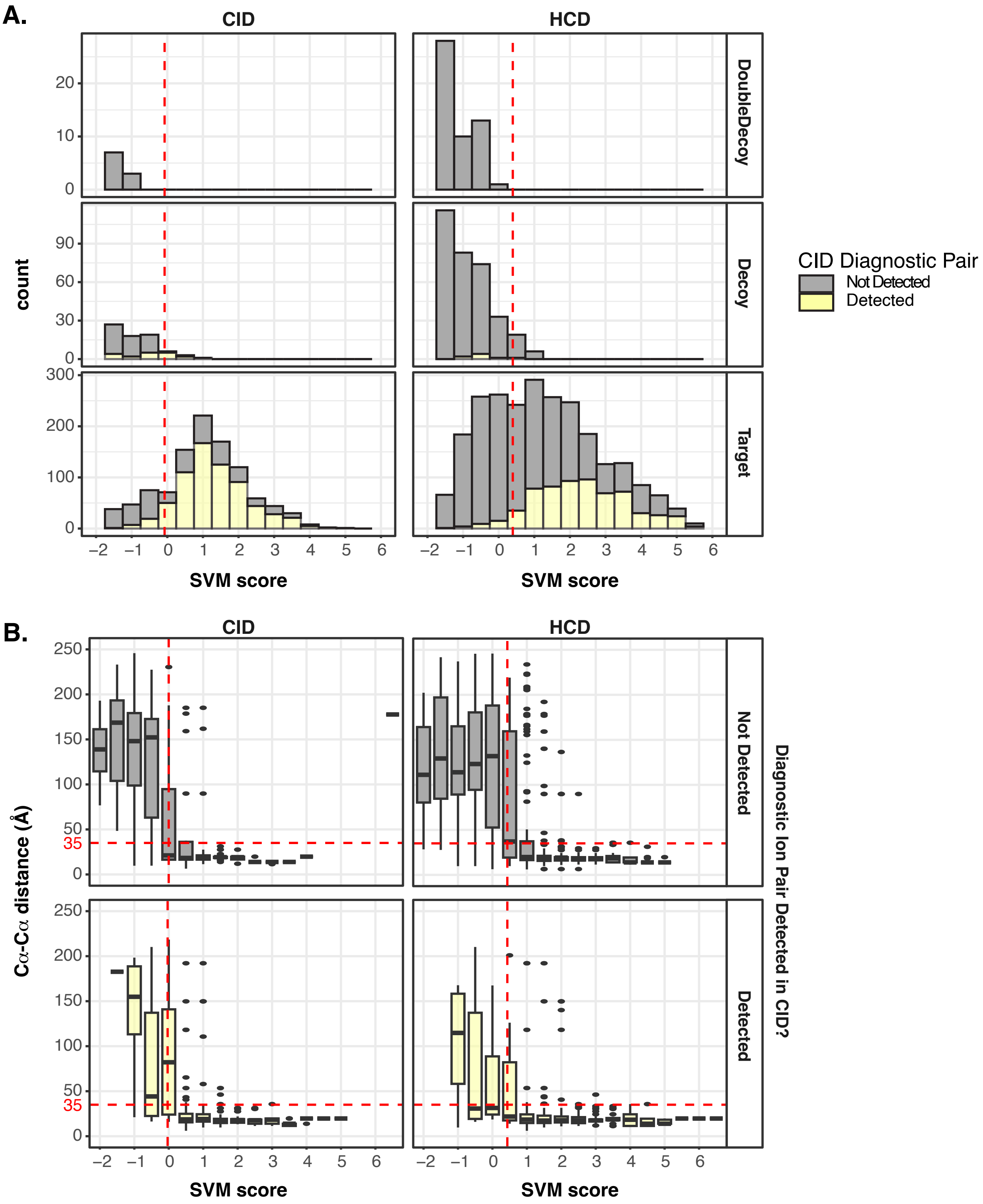

MS2 CSMs

DSSO, rabbit Ribosome

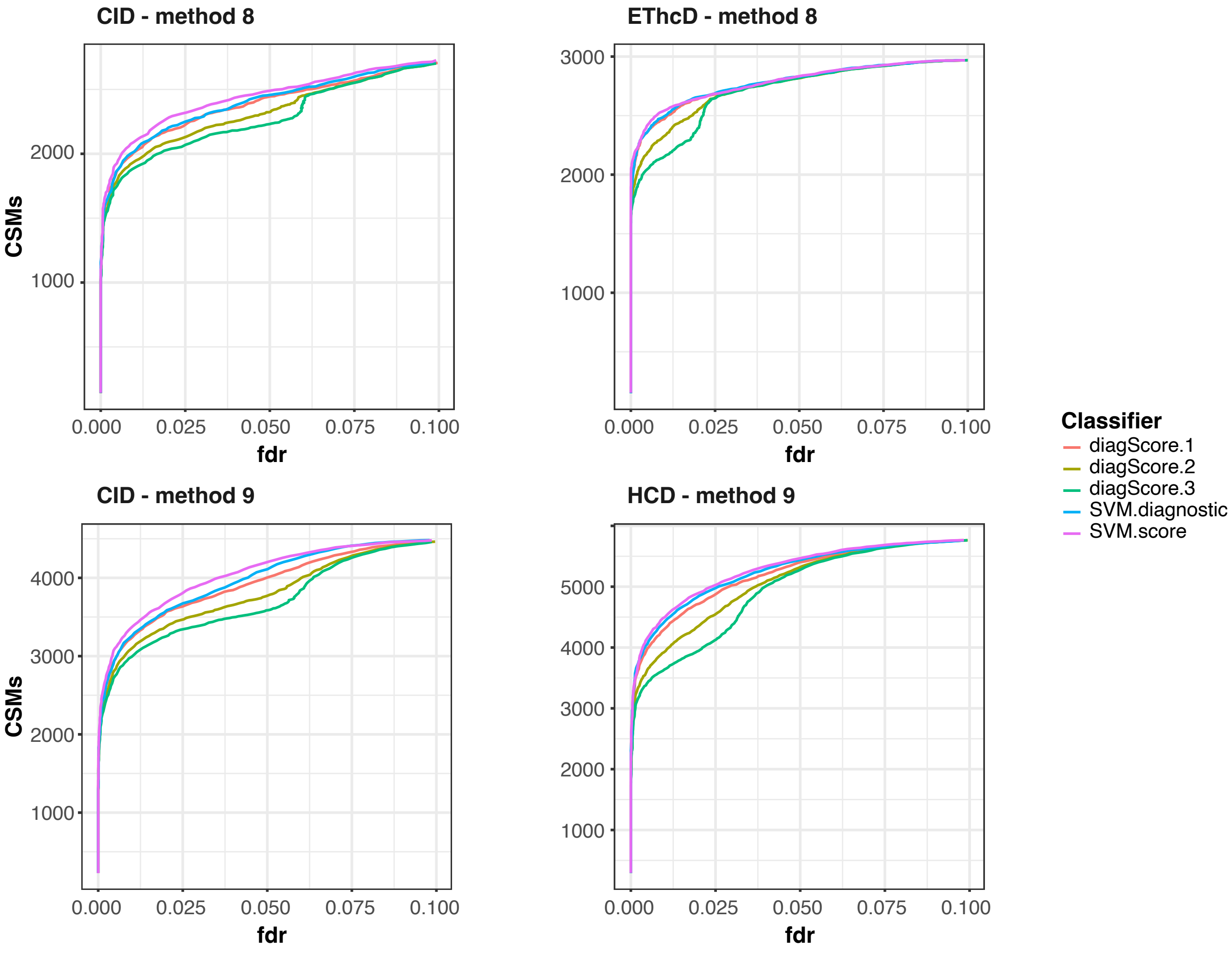


**Supplemental Figure 12.**

The detection of diagnostic DSSO ion pairs does not improve scoring of crosslinked spectral matches in MS2. The performance of different scoring functions was evaluated on the DSSO crosslinked Rabbit Ribosome data acquired by sequential CID-EThcD (top panels) or sequential CID-HCD (bottom panels) by plotting the number of CSMs classfied at different FDR thresholds. SVM.score, the standard Touchstone scoring function which does not consider diagnostic pair detection, classified more spectra at all FDR thresholds than scoring functions which incorporated diagnostic ion pairs. SVM.diagnostic is trained on the same features as SVM.score plus diagnostic ion pair detection. diagScore.x - are scoring functions created by adding 1 to 3 points to SVM.score when a diagnostic pair was detected in the CID spectrum (eg, HCD and EThcD scans detect diagnostic ion pairs in the corresponding CID of the same precursor ion).

#### MS2 CSMs

DSSO, rabbit Ribosome

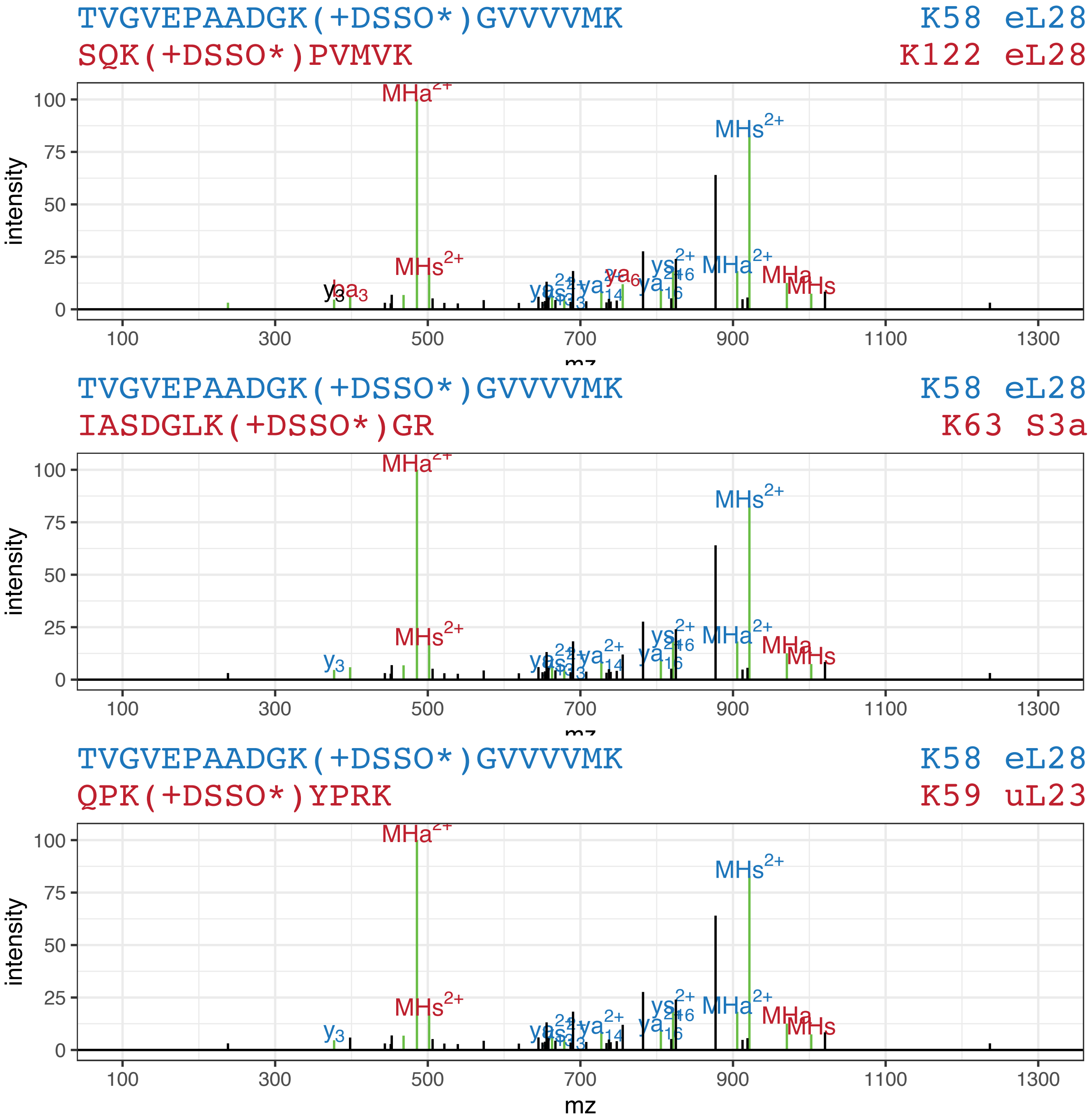


**Supplemental Figure 13.**
The presence of diagnostic ion pairs has low utility in discriminating correct from incorrect crosslink matches. The same spectrum matches three distinct crosslinked peptide pairs. The best scoring peptide (peptide 1) is the same in all cases but the matches differ in the peptide matching to the second peptide. In all cases, the MHa and MHs diagnostic ion pairs are matched and the correct intraprotein hit on Ribosomal subunit L28 is distinguished by more backbone ion matches.

**Supplemental Figure 14.**
Precursor ion m/z distribution (top) and z distribution (bottom) from HEK293 DSSO crosslinking with different methods

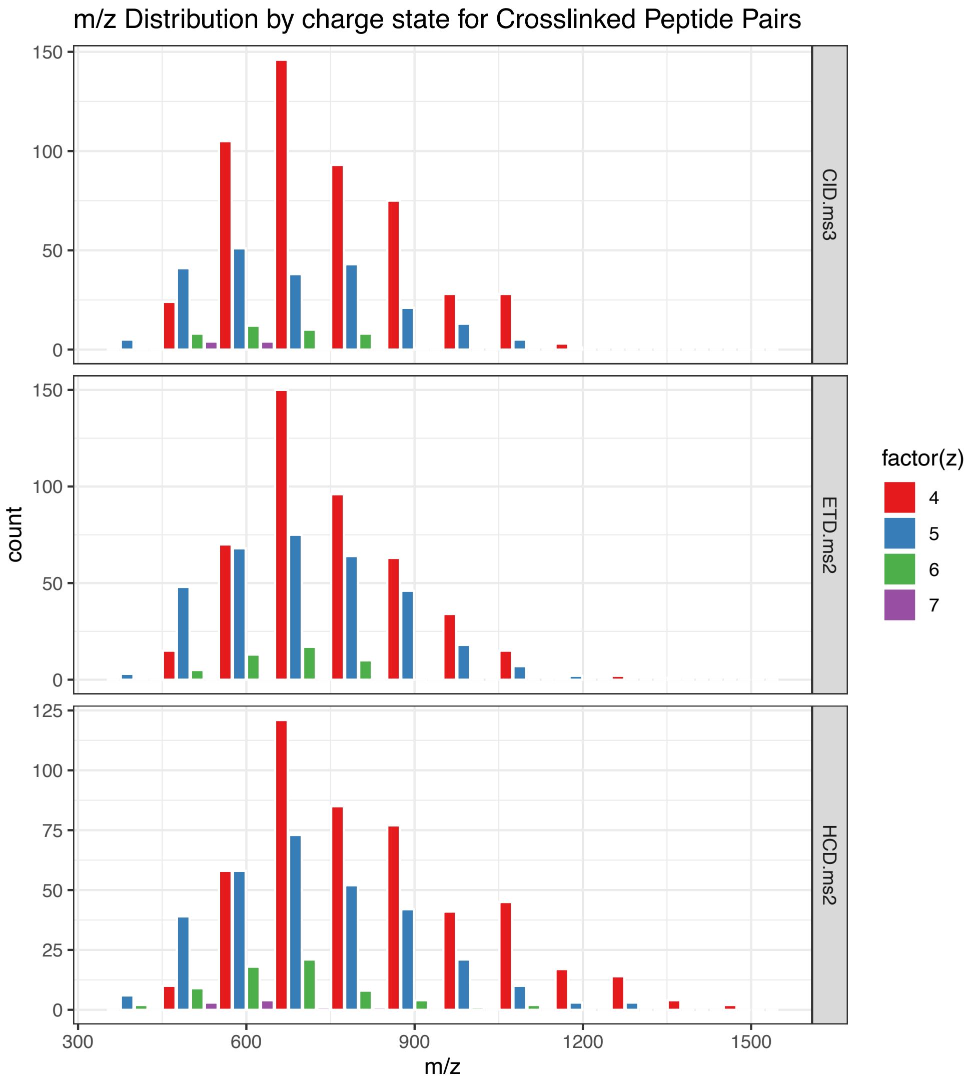


Fraction of Spectra Identified, by charge-state, technique

| z | CID | ETD.ms2 | HCD.ms2 |
|---|---|---|---|
| 4 | 6.28 | 6.25 | 6.04 |
| 5 | 7.19 | 9.23 | 8.17 |
| 6 | 5.09 | 6.61 | 6.70 |
| 7 | 2.74 | 3.47 | 4.04 |
| 8 | 0.70 | 1.03 | 1.42 |
| Total | 6.37 | 6.98 | 6.61 |

**Supplemental Figure 15.**
Crosslinked Peptide length distribution from HEK293 DSSO crosslinking analysis acquired by different MS methods

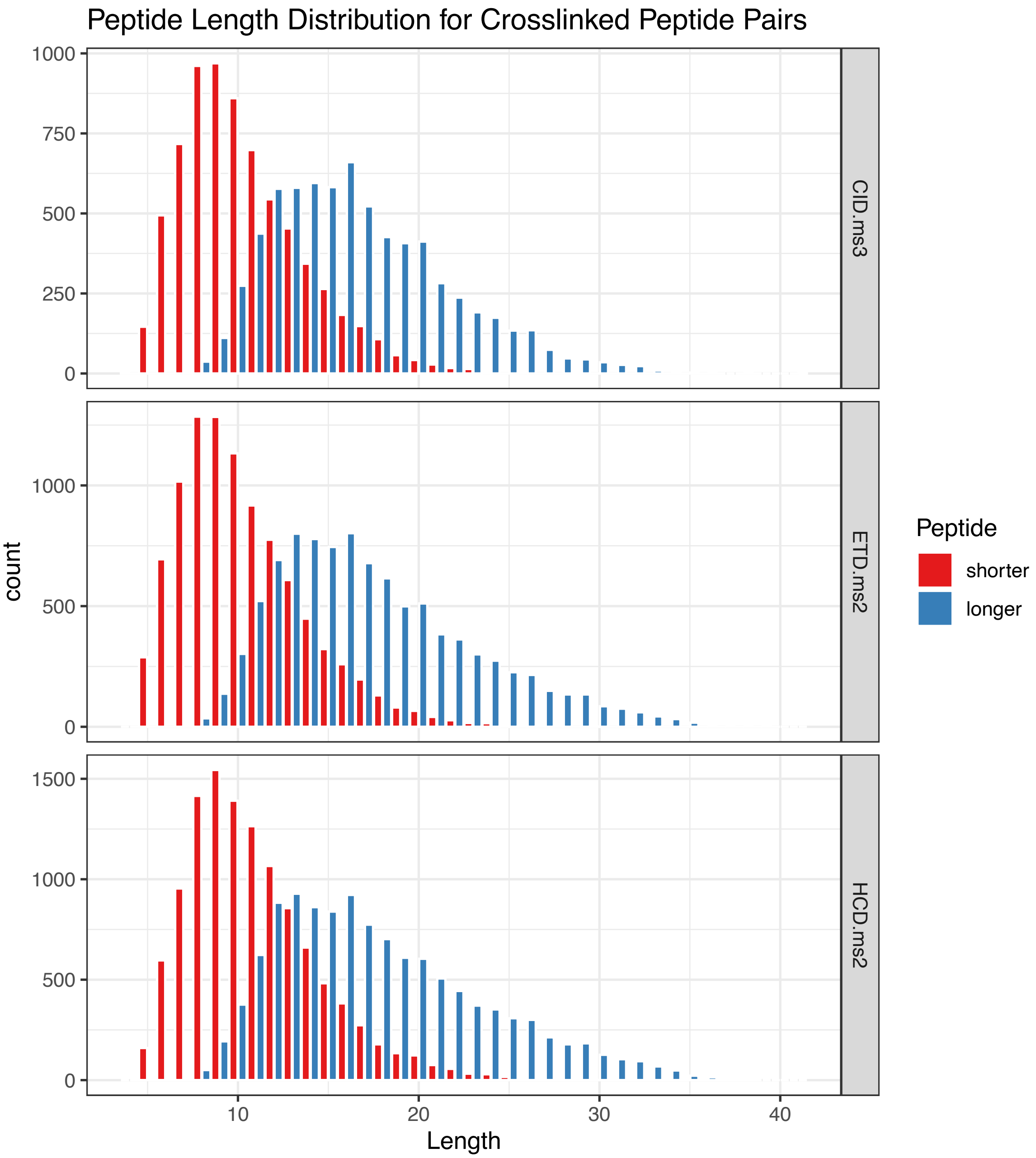
